# Physiological Cryptic Last Exon Splicing Generates Regulated and Functional Transcript Isoforms Across Vertebrates

**DOI:** 10.64898/2026.01.30.702498

**Authors:** Matthew P. Bostock, Karla L. Gonzalez, Colin Lentjes, Samina Uddin, Lourdes O. Almagro, Fursham Hamid, Corinne Houart

## Abstract

Cryptic last exons (CLEs) are unannotated terminal exons that arise from intronic sequence and have predominantly been associated with RNA-binding-protein dysfunction and neurodegenerative disease. Whether CLEs also contribute to physiological gene regulation remained unclear. Here, we identify thousands of CLE-containing transcripts across wild-type zebrafish, mouse and human transcriptomes. CLE sequences are poorly conserved, yet CLE occurrence recurs in orthologous genes and, less frequently, orthologous introns, suggesting that gene architecture creates recurrent opportunities for CLE formation. A subset of CLEs is developmentally regulated and CLE usage is unusually sensitive to perturbation of neural RNA-binding proteins. CLE-containing transcripts associate with ribosomes, and endogenous tagging demonstrates that *ephA4b-CLE* produces a stable truncated protein during normal development. Loss of *ephA4b-CLE* impairs retinal ganglion cell axon growth, whereas disruption of *GAN-CLE* increases motor-axon extension and terminal branching. Moreover, independently arising human and zebrafish *EphA4* CLEs converge on similar predicted ephrin-binding ectodomains despite lacking sequence conservation. These findings establish CLEs as a physiological source of transcript diversity within which individual isoforms can acquire regulated and gene-specific developmental functions.

## Introduction

The vertebrate nervous system exhibits extraordinary cellular diversity generated by developmental programmes that specify neuronal identity and connectivity across space and time. Achieving this complexity required transcript-specific regulation beyond transcription alone. Alternative splicing provides this regulation by generating distinct mRNA isoforms that diversify the proteome and specify neuronal identity^1–6^. Multiple modes of alternative splicing have been described, including exon skipping, mutually exclusive exons, alternative 5’/3’ splice site, intron retention, microexons, poison exons, and alternative first/last exons (AFE/ALE)^7–17^.

Although canonical splicing modes are well characterised, high resolution transcriptomic analyses have revealed novel, atypical events, the biological significance of which remains unclear. One such class is cryptic exons, unannotated exonic sequences occurring in intronic regions that are typically repressed by RNA binding proteins (RBPs) like TAR DNA-binding protein 43 (TDP-43) or Splicing Factor Proline and Glutamine Rich (SFPQ)^18–21^. Their inclusion may disrupt or modify canonical mRNAs, often leading to reduced levels of protein^21,22^.

A specialised subset of cryptic exons, called cryptic last exons (CLEs), arises when intronic splice acceptor sites are coupled to premature stop codons and polyadenylation sites^19,22^. Unlike most cryptic exons, CLE inclusion results in premature transcript termination without eliminating transcript production, generating shortened mRNAs predicted to encode truncated proteins^19,23^. CLEs were the most highly expressed alternative splicing event in zebrafish *sfpq* null mutant, SFPQ/TDP-43-deficient mouse cortex and human ALS patient-derived iPSCs and were responsible for developmental abnormalities such as brain boundary formation defects^19^. These findings established the prevailing view that CLEs are pathological splicing errors that are normally suppressed in healthy tissue.

However, several observations suggest they may not be purely pathological. CLEs display characteristics of stable RNAs like defined branch points and functional polyadenylation sites. Moreover, CLE-containing transcripts have been detected at low levels in wild-type RNA sequencing (RNA-seq) datasets and in control zebrafish samples through RT-PCR, observations that have been generally attributed to accidental by-products of splicing^19,21^. Whether these transcripts reflect accidental splicing noise or instead represent an unrecognised mode of transcript regulation is unresolved.

Here, we systematically characterise CLE usage across physiological zebrafish, mouse and human transcriptomes. We show that CLEs are predominantly low-usage isoforms but that a subset is developmentally regulated and unusually responsive to changes in neural RBP state. Although CLE sequences show little evolutionary conservation, CLE formation recurs in orthologous genes and intronic contexts, suggesting recurrent acquisition within permissive gene architectures. We further show that CLE transcripts can associate with ribosomes and directly demonstrate endogenous production of an EphA4b-CLE protein. Functional disruption of *ephA4b-CLE* and *GAN-CLE* produces distinct neuronal developmental phenotypes, while independently arising human and zebrafish *EphA4* CLEs converge on similar predicted ligand-binding ectodomains. Together, these findings establish CLEs as a physiological class of alternative transcript termination events capable of generating gene-specific regulatory and functional outputs.

## Results

### CLEs are expressed in a diverse set of zebrafish, mouse, and human wildtype cell types

If CLEs are not solely pathological products of splicing-factor dysfunction, CLE-containing transcripts should also be detectable under physiological conditions. To determine whether physiological CLE usage extends across vertebrates, we performed a large-scale analysis of publicly available wild-type RNA-seq datasets from zebrafish, mouse, and human across multiple developmental stages (Supplementary Datasheet 1), using our transcriptome-assembly pipeline to identify *de novo* CLE-containing transcripts (Figure 1A)^28^.

**Figure 1:**
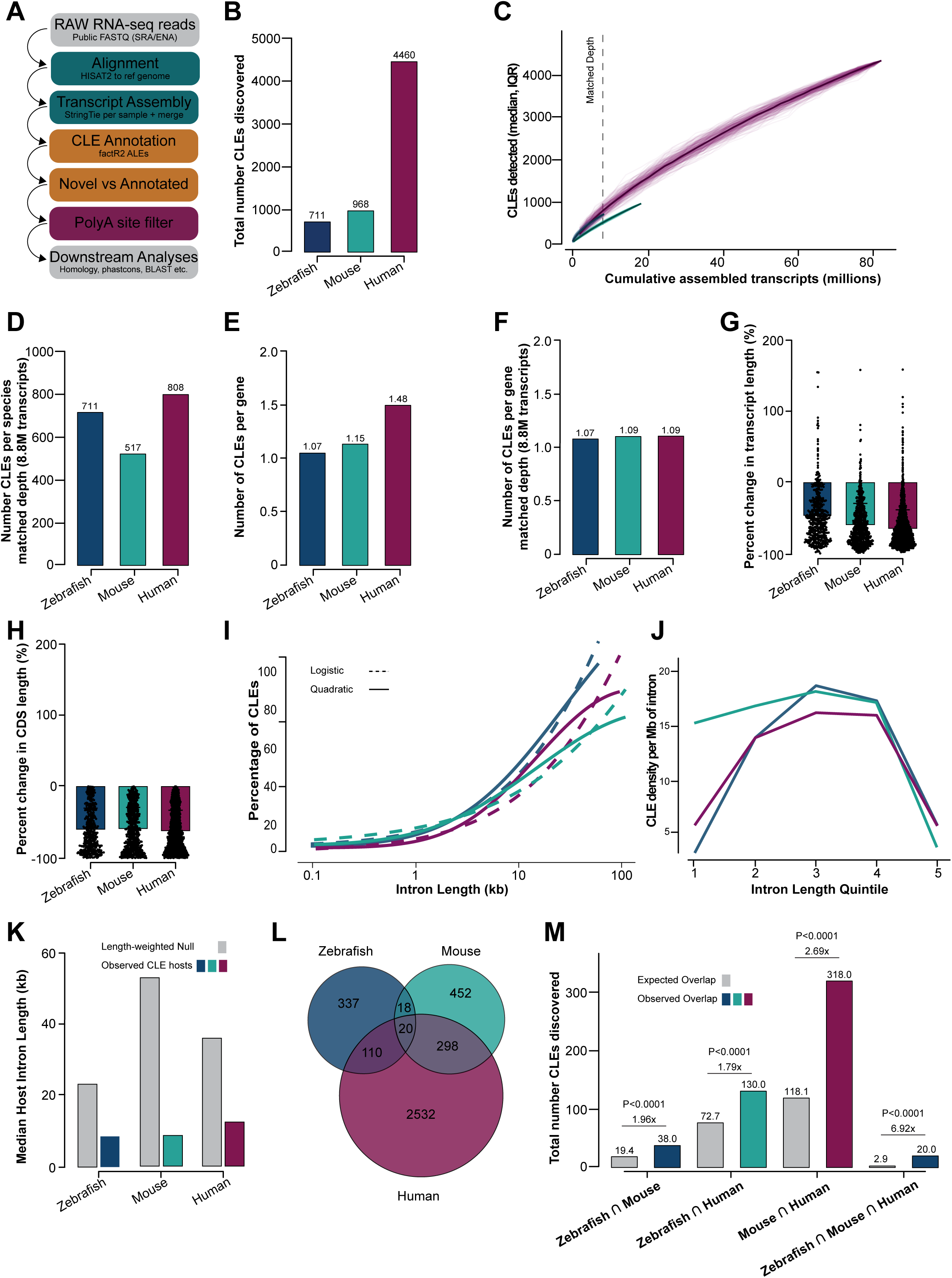
Cryptic last exons are widely detected across physiological vertebrate transcriptomes and recur in orthologous genes. **(A)** Schematic of the CLE-discovery pipeline. Public physiological RNA-seq datasets were aligned using HISAT2, assembled using StringTie, and analysed using factR2 to identify novel alternative last exons, followed by polyadenylation-signal and splice-junction-support filtering. **(B)** Total numbers of CLEs identified in zebrafish (711), mouse (968), and human (4,460). **(C)** Rarefaction analysis showing cumulative CLE discovery as a function of cumulative assembled-transcript depth. Bold lines show median CLE recovery and transparent lines show the 200 random sample orderings. Dashed line indicates the maximum assembled-transcript depth shared by all three species (∼8.8 million transcripts). **(D)** Number of CLEs recovered at matched assembled-transcript depth: zebrafish, 711; mouse, 517; human, 808. **(E–F)** Mean number of CLEs detected per CLE-host gene using the complete datasets (E) or following matching to equivalent assembled-transcript depth (F). **(G–H)** Percentage change in mature transcript length (G) and predicted coding-sequence length (H) of CLE-containing transcripts relative to the corresponding canonical transcript. Negative values indicate truncation. **(I)** Relationship between intron length and probability of CLE occurrence. Dashed lines show predictions from the linear logistic model and solid lines from the model containing a quadratic intron-length term. Inclusion of the quadratic term significantly improved model fit in zebrafish, mouse and human (likelihood-ratio tests, P=1.3×10⁻⁵, 1.8×10⁻¹² and 6.2×10⁻⁷⁹, respectively). **(J)** Density of CLE-containing introns per Mb of intronic sequence across species-specific intron-length quintiles. **(K)** Median length of observed CLE-host introns compared with the median expected under a length-weighted null model in which introns were sampled in proportion to available intronic sequence. Observed CLE-host introns fell below all 1,000 null draws in each species (one-sided empirical randomisation test, P=0.001). **(L)** Overlap of CLE-hosting orthologous genes across zebrafish, mouse, and human. (M) Observed numbers of shared CLE-hosting orthologues compared with the expectation from 10,000 random gene-set permutations. Observed overlaps were enriched 1.96-fold for zebrafish–mouse, 1.79-fold for zebrafish–human, 2.69-fold for mouse–human, and 6.92-fold across all three species (all P<0.0001). **(M)** Observed numbers of shared CLE-hosting orthologues compared with expectation from 10,000 random gene-set permutations. Observed overlaps were enriched 1.96-fold for zebrafish–mouse, 1.79-fold for zebrafish–human, 2.69-fold for mouse–human and 6.92-fold across all three species (empirical permutation tests, all P<1×10⁻⁴; Benjamini–Hochberg FDR<1×10⁻⁴).

We identified 711 CLE events in 652 genes in zebrafish, 968 in 833 genes in mouse, and 4,460 in 2,960 genes in human (Figure 1B; Supplementary Datasheet 2). All retained CLEs were supported by a StringTie-assembled transcript in at least one individual RNA-seq sample, with direct splice-junction support for the CLE-containing isoform and contained a predicted open reading frame terminating within the CLE. However, the amount of RNA-seq data available differed substantially between species, comprising 168 zebrafish, 279 mouse and 3,088 human samples (Supplementary Datasheet 1). We therefore asked whether the approximately six-fold larger human CLE catalogue reflected greater sampling effort rather than a biological increase in CLE usage.

To account for this difference, we performed a rarefaction analysis in which biological samples were repeatedly added in random order and the cumulative number of unique CLEs was recorded against the cumulative number of assembled transcripts. Repeating this process 200 times generated an accumulation curve for each species showing how CLE discovery increased as progressively more transcriptomic data were sampled (Figure 1C). As expected, additional sampling continued to reveal new CLEs. We compared the three species at the same cumulative transcript depth, using 8.88 million assembled transcripts as the maximum depth shared by all three datasets (Figure 1C). At this matched depth, 711 CLEs were detected in zebrafish, 517 in mouse and 808 in human, compared with the raw totals of 711, 968 and 4,460, respectively (Figure 1B). Thus, the large excess of CLEs in the complete human catalogue largely disappeared when sampling effort was matched, indicating that raw cross-species differences in CLE number predominantly reflect the amount of transcriptomic data available rather than a lineage-specific expansion of CLE usage.

A similar effect was apparent when considering the number of CLEs detected per CLE-containing gene. In the complete datasets, the mean number of CLEs per gene was 1.07 in zebrafish, 1.16 in mouse and 1.48 in human (Figure 1D). At matched transcript depth these values converged to 1.07, 1.09 and 1.09, respectively (Figure 1E), indicating that the apparent increase in multiple CLEs per human gene was also largely attributable to greater sampling. Overall, CLE discovery frequencies were similar across species after matching transcriptomic sampling depth.

Having established that CLE-containing transcripts occur physiologically, we next asked to what extent CLEs contribute to host-gene transcript usage. CLE inclusion was quantified from junction reads as the proportion of splice junctions from the upstream donor terminating at the CLE relative to the CLE and competing full-length junctions [CLE/(CLE+FL)], considering CLEs supported by at least 10 informative junction reads. CLEs predominantly represented non-dominant transcript isoforms, with median inclusion levels of 0.05 in zebrafish, 0.01 in mouse and 0.02 in human (Supplementary Figure 1A). Consistent with this, 83.7% of zebrafish, 97.1% of mouse and 92.8% of human CLEs were used as the non-dominant isoform (PSI<0.5), with PSI distributions significantly below 0.5 in all three species (Wilcoxon signed-rank test, all P<1×10⁻⁵⁰). More specifically, 69.0% of zebrafish, 94.8% of mouse and 84.9% of human CLEs had inclusion below 20%, whereas only 10.3%, 1.3% and 3.3%, respectively, reached PSI values above 0.8 (Supplementary Figure 1A). Thus, physiological CLE inclusion is generally low across vertebrates, but spans a broad range of inclusion frequencies, with a minority of CLEs becoming major transcript isoforms. Because many of the source datasets were generated from bulk tissues, low aggregate CLE inclusion may reflect either uniformly low usage across cells or higher inclusion within restricted cell types or cell states.

Since CLEs introduce a premature terminal exon and polyadenylation site, we next asked how CLE inclusion alters mature transcript structure and predicted protein-coding potential. For each CLE, we compared the CLE-terminated isoform with the Ensembl canonical transcript of its host gene and calculated the percentage change in mature transcript length (Figure 1G). CLE inclusion substantially shortened both features across all three species, with median reductions in mature transcript length of 55.4% in zebrafish, 66.3% in mouse and 70.7% in human (Figure 1G). Using factR2 open reading frame predictions, we similarly calculated the change in predicted protein length (Figure 1H). Predicted protein length was similarly reduced by a median of 68.8% in zebrafish, 64.8% in mouse and 68.1% in human (Figure 1H). Although a small number of CLE-containing transcripts were longer overall than their corresponding canonical transcripts because of an extended CLE-derived 3′ untranslated region, translation terminated earlier in every case, resulting in a shorter predicted protein. CLE inclusion therefore predominantly shortens mature transcripts and consistently truncates the predicted protein product.

Pathogenic CLEs were reported to appear in long introns^19^, thus, we assessed whether this was the case for the CLEs discovered in this study. We ranked introns of each CLE-containing gene by descending size (rank 1 being the longest intron) and determine which intron ranks CLEs occupy. CLEs were frequently located within the longest intron of their host genes (87.42% in zebrafish; 79.01% in mouse, 56.98% in human; Supplementary Figure 1C). Because longer introns provide more sequence in which a CLE could arise, we asked whether the association between CLE occurrence and intron length could be explained solely by available intronic sequence. We first modelled the probability that an intron contained a CLE as a function of its length using logistic regression. A model in which CLE probability changed monotonically with log intron length showed a significant positive association in all three species (log-odds per log₁₀-length: zebrafish +1.68, mouse +1.21, human +1.54; all P < 1×10⁻^95^, Wald z-tests). We then added a quadratic length term, allowing the association to rise and subsequently weaken rather than increase without bound. This non-linear model significantly improved fit in zebrafish, mouse and human (likelihood-ratio test P = 1.3×10⁻⁵, 1.8×10⁻¹² and 6.2×10⁻⁷⁹, respectively), with a significant negative quadratic term in each species (loglen² = −0.44, −0.46 and −0.77; all P ≤ 6.5×10⁻⁵, Wald z-tests, Supplementary Datasheet 3), indicating that the positive association with length weakens and ultimately reverses in the longest introns (Figure 1I). Consistent with this, CLE density across intron-length quintiles was non-monotonic, increasing across short-to-intermediate introns and then falling sharply in the longest quintile (Figure 1J). CLE density peaked in the third quintile (≈1.2–2.5 kb in zebrafish, ≈1.5–3.1 kb in mouse, ≈1.7–3.6 kb in human) and collapsed in the longest quintile (>5.5 kb, >8.5 kb and >10 kb, respectively): in mouse, density was 15.0, 16.5, 17.8, 16.8 and 3.7 CLEs per Mb across increasing length quintiles, with similar patterns in zebrafish (6.0, 16.4, 21.2, 19.8, 8.5) and human (5.0, 13.0, 15.3, 15.0, 5.0) (Figure 1J). Finally, we compared observed CLE-host intron lengths with a length-weighted null representing the expectation if CLE occurrence were proportional simply to available intronic sequence (Figure 1K). Observed CLE-host introns were substantially shorter than this expectation in every species, with median observed versus expected lengths of 8,474 versus 23,236 bp in zebrafish, 8,653 versus 52,892 bp in mouse and 12,362 versus 36,044 bp in human; the observed median fell below all 1,000 length-weighted null draws in each species (one-sided empirical randomisation test, P < 0.001; Figure 1K, Supplementary Datasheet 3). Intron rank was independently associated with CLE occurrence, with earlier introns more likely to host a CLE after accounting for intron length (rank log-odds: zebrafish −0.042, mouse −0.045, human −0.029; all P < 1×10⁻⁸, Wald z-tests, Supplementary Datasheet 3). Together, these analyses show that CLE occurrence is not explained simply by greater sequence availability in long introns, but instead reflects a shared preference for introns of intermediate length and earlier position across vertebrate species.

We next asked if CLE occurrence recurs in orthologous genes across vertebrate species. To do this, CLE-hosting genes from each species were mapped onto a common set of human orthologue symbols, and the observed overlaps were compared with null distributions generated from 10,000 random gene-set permutations that preserved the number of CLE-hosting genes identified in each species. CLE-containing orthologues overlapped significantly more frequently than expected in all pairwise comparisons. Zebrafish and mouse shared 38 CLE-hosting orthologues compared with 19.4 expected under the null, representing a 1.96-fold enrichment. Zebrafish and human shared 130 compared with 72.7 expected, a 1.79-fold enrichment, while mouse and human shared 318 compared with 118.1 expected, a 2.69-fold enrichment (Figure 1L-M). The recurrence was particularly pronounced across all three species, with 20 orthologous genes containing a CLE in zebrafish, mouse and human compared with only 2.9 expected by chance, representing a 6.92-fold enrichment. In each comparison, the observed overlap exceeded that obtained in all 10,000 permutations (P<1×10⁻⁴; Benjamini-Hochberg FDR<1×10⁻⁴).

We next asked whether recurrence within the same orthologous genes reflected conservation of the individual CLE events at equivalent positions, that is, whether CLEs in shared host genes occupy orthologous introns across species. We defined orthologous introns as being flanked by two conserved exons. For every CLE in a shared gene, we projected its two flanking exons into the partner species genome by liftOver and asked whether the partner species CLE fell within the resulting orthologous intron. A substantial fraction of testable shared-gene CLEs occupied the orthologous intron, and this fraction decreased with evolutionary distance, from 28.5% of testable pairs in the mouse-human comparison (86/302) to 13.0% for zebrafish-human (9/69) and 9.1% for zebrafish-mouse (2/22). The signal was specific: randomly re-pairing host genes abolished it (permutation null approximately 0%; Supplementary Datasheet 4), and 91 of 97 orthologous-intron calls were independently confirmed by reciprocal liftOver of their flanking exons (Supplementary Datasheet 4). No CLE occupied the orthologous intron across all three species simultaneously (0 of 19 genes shared by all three), and positional conservation among these genes was confined to individual species pairs (7 of 19 in at least one pair), predominantly the closely related mouse-human comparison. Thus, while the CLE sequences themselves are not conserved, their intronic positions are partially retained between more closely related species, indicating that specific intronic contexts are recurrently predisposed to CLE formation rather than that ancestral CLEs are faithfully maintained across vertebrate evolution.

Together, these results distinguish gene-level recurrence from conservation of individual cryptic exons. Some orthologous genes are repeatedly associated with CLE formation across vertebrate species, but the CLEs themselves generally arise at different intronic positions. This pattern is therefore more consistent with recurrent or independent acquisition of cryptic terminal exons within susceptible host genes than with evolutionary maintenance of individual CLE events.

### <u>CLE sequences are evolutionarily non-conserved, transposon-rich, and</u> <u>lineage restricted.</u>

Given that CLE formation recurs in orthologous genes and, less frequently, orthologous introns, we next asked whether CLE sequences themselves show evidence of evolutionary conservation. We compared phastCons scores (Figure 2A) of CLEs against alternative last exons (ALEs), size-matched intronic sequences, upstream-exon controls, and constitutive exons all gene-matched to the CLE host across the three species. CLEs were among the least conserved sequence classes in every species (median phastCons 0.000 in zebrafish, 0.049 in mouse, 0.038 in human, Figure 2B, Supplementary Datasheet 5). Their conservation was negligibly different from that of size-matched intronic sequences (raw median differences 0.000, −0.006, −0.008 in zebrafish, mouse, and human, Supplementary Datasheet 5) and far below ALEs (raw median differences −0.467, −0.175, - Supplementary Datasheet 5), upstream exons (raw median differences −0.837, −0.713, −0.686), and constitutive exons (raw median differences ∼-0.82 to −0.89, Supplementary Datasheet 5). Pairwise rank-based comparisons gave the same results, with CLEs indistinguishable or only minimally different from intronic sequences and strongly separated from all exonic controls (Hodges-Lehmann differences and adjusted p-values in Supplementary Datasheet 5). Thus, unlike bona fide ALEs, CLE sequences show essentially no evolutionary constraint (Figure 2B, Supplementary Datasheet 5).

**Figure 2:**
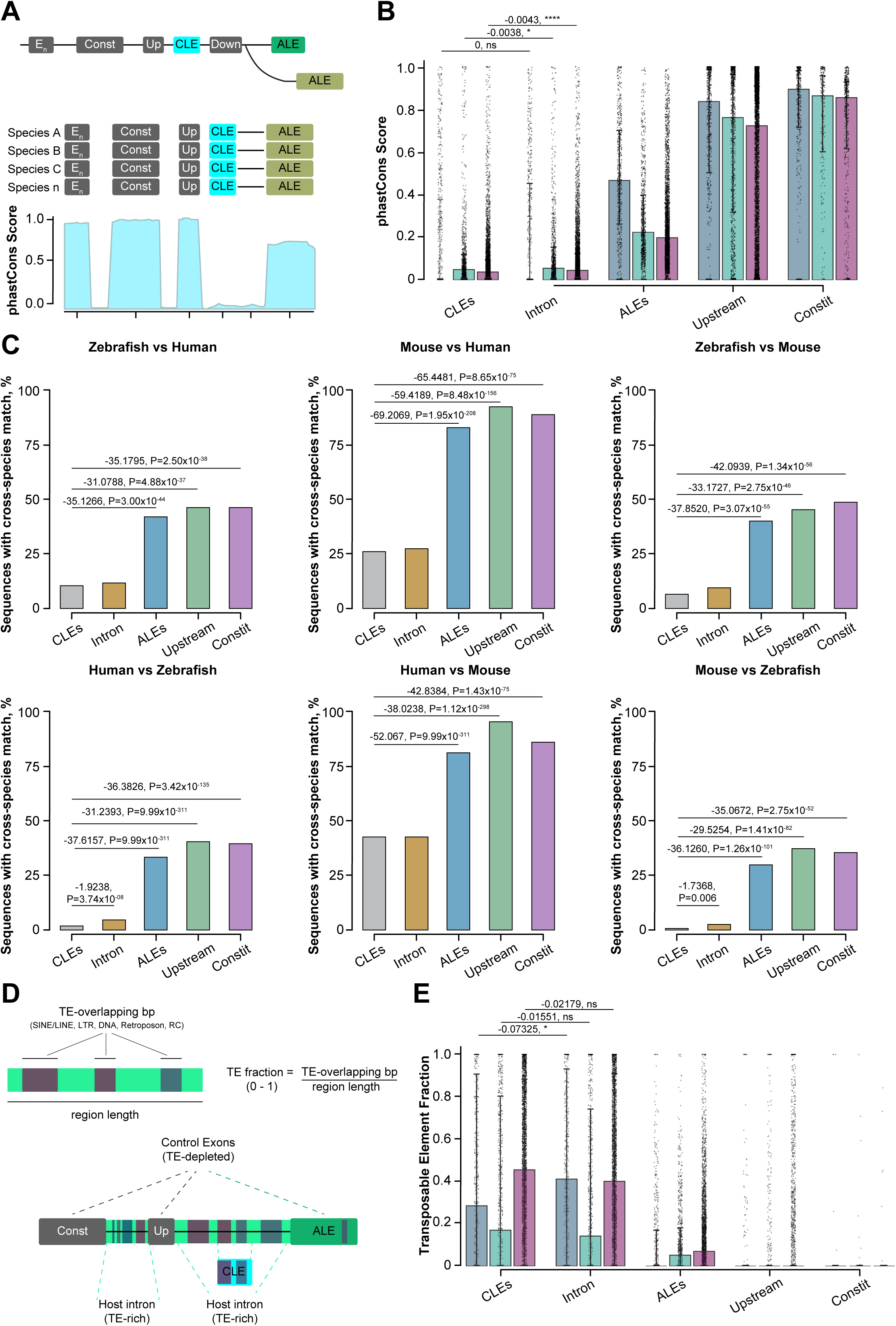
CLE sequences are poorly conserved and retain the sequence characteristics of their host introns. **(A)** Schematic of the gene-matched sequence classes used for evolutionary comparisons. CLEs were compared with size-matched intronic regions, conventional alternative last exons (ALEs), the nearest upstream annotated exon, and constitutive internal exons from CLE-host genes. Mean phastCons conservation was calculated across each region. **(B)** phastCons conservation scores for CLEs and control regions in zebrafish, mouse, and human. CLEs showed conservation scores similar to size-matched intronic sequence and substantially lower than ALEs, upstream exons, and constitutive exons. Median CLE phastCons scores were 0.000 in zebrafish, 0.049 in mouse, and 0.038 in human. Pairwise comparisons were performed using two-sided Wilcoxon rank-sum tests with Benjamini–Hochberg correction. **(C)** Percentage of CLE and control sequences producing a cross-species genomic BLAST match in each of the six directional zebrafish, mouse, and human comparisons. CLE sequences showed substantially lower cross-species hit rates than ALEs, upstream exons, and constitutive exons and were similar to intronic controls. Statistical comparisons shown are Fisher’s exact tests with Benjamini–Hochberg correction. **(D)** Schematic of transposable-element (TE) analysis. TE fraction was calculated as the proportion of each region overlapping SINE, LINE, LTR, DNA, Retroposon, or rolling-circle RepeatMasker annotations. **(E)** TE fraction of CLEs and gene-matched intronic, ALE, upstream-exon, and constitutive-exon controls in zebrafish, mouse, and human. CLEs were strongly enriched for TE sequence relative to annotated exonic controls and showed TE content similar to their surrounding intronic sequence. Statistical comparisons were performed using two-sided Wilcoxon rank-sum tests with Benjamini–Hochberg correction; ns, not significant; *P<0.05; ****P<0.0001.

PhastCons measures conservation at each aligned position, therefore CLEs would only align if they occur at the same locus across species. Homologous CLE sequence residing at a different genomic location in another species would be missed by this positional measure, which could explain the low phastCons scores of CLEs (Figure 2B). To test for such position-independent homology, we performed pairwise BLAST searches of each CLE sequence across zebrafish, mouse, and human. Consistent with their low conservation, CLE sequences found cross-species genomic matches far less often than ALEs (for example, 22.4% of CLEs versus 91.6% of ALEs in the mouse-to-human comparison; odds ratio 0.027, p<4.07×10^-209^, Figure 2C, Supplementary Datasheet 5). This trend occurred across all six pairwise species comparisons: CLE hit rates were 4-to 40-fold lower than those of ALEs (odds ratios 0.017 to 0.135; all p < 2.00×10^-44^, Supplementary Datasheet 5) and, in every comparison, were either not significantly or minimally different from those of intronic control sequence (for example 22.4% vs 24.0% for mouse-to-human, p = 0.41, Supplementary Datasheet 5), while remaining far below constitutive and upstream exons. Together, phastCons and BLAST analyses showed that CLEs are as conserved as intronic sequences and much less than ALEs, upstream exons and constitutive exons.

CLE sequences so far resembled intronic regions rather than ALEs, upstream exonic controls, or constitutive exons. To further probe this intronic character, we quantified the occurrence of transposable elements (TEs) in CLEs and controls using RepeatMasker annotations from the UCSC Genome Browser (Figure 2E). TE enrichment is a feature of introns and tend to be absent in exons^29,30^. CLEs were markedly TE-enriched relative to ALEs: TEs covered a median of 29% of CLE bodies in zebrafish, 17% in mouse, and 46% in human, compared with close to zero for ALE bodies (raw median differences −28.5%, −11.8%, and −38.4%; Figure 2F, Supplementary Datasheet 5). Pairwise rank-based comparisons confirmed this enrichment (Cliff’s Δ 0.36 in zebrafish, 0.20 in mouse, and 0.40 in human; all p<1×10^-14^; Supplementary Datasheet 5). The enrichment held within individual genes: in genes hosting both a CLE and an ALE, CLE bodies were substantially more TE-rich than the gene’s own ALEs (median within-gene difference 0.146 in zebrafish, 0.101 in mouse, and 0.329 in human; paired Wilcoxon p from 6×10^-38^ to 2×10^-272^ across 578, 833, and 2,960 paired genes; Supplementary Datasheet 5), showing that the TE enrichment of CLEs is not a consequence of gene-level composition differences between the CLE- and ALE-hosting gene sets. In contrast to this strong enrichment over exons, CLE TE content closely matched that of intronic sequence (median 41% in zebrafish, 14% in mouse, and 40% in human; differences non-significant except in zebrafish, p < 0.005; Figure 2F, Supplementary Datasheet 5), consistent with CLEs deriving from and retaining the repeat character of the introns in which they reside.

Lastly, we hypothesised that CLEs occurring in orthologous genes across species may be more conserved. To test this, we compared phastCons scores of CLEs whose host gene carried a CLE in two or more species (“shared”, n = 1,277) against species-specific CLEs (“private”, n = 4,692). Shared CLEs were no more conserved than private ones (median phastCons 0.039 versus 0.038, pooled, Supplementary Figure 2A; not significant in any species, Supplementary Figure 2B, Supplementary Datasheet 5). This indicated that the recurrence of CLEs in orthologous genes is a property of the host gene and not due to a conserved underlying sequence. Shared CLEs did, however, differ in their relationship to TEs. Shared CLEs contained less TE sequences than private CLEs (pooled median TE fraction 0.21 versus 0.44; Cliff’s Δ −0.14, p = 3×10-14, Supplementary Figure 2C, Supplementary Datasheet 5). This trend was significant in mouse and human and trending in the same direction in zebrafish (Supplementary Figure 2D, Supplementary Datasheet 5). This lower TE content reflected a shift in their presence or absence rather than their extent: shared CLEs were markedly more likely to be entirely TE-free (41.6% versus 30.2% TE-free, pooled; odds ratio 1.65, p = 4×10-14, Supplementary Figure 2E), a difference seen in all three species (zebrafish 47.9% versus 37.7%; mouse 52.5% versus 37.9%; human 34.3% versus 28.0%, Supplementary Figure 2F, Supplementary Datasheet 5). Thus, while CLEs as a class are TE-rich, CLEs recurring in orthologous host genes are disproportionately TE-poor or TE-free. This may reflect more frequent emergence from TE-independent intronic sequence, or alternatively evolutionary divergence of ancestral TE-derived sequence beyond current repeat annotation; the present data do not distinguish between these possibilities.

CLEs occupying orthologous introns were indistinguishable from other shared-gene CLEs in sequence conservation, cross-species alignment, and TE content (phastCons, BLAST, and TE fraction all non-significant against a shared-CLE background; Supplementary Datasheet 5). Positional recurrence is therefore not associated with any distinct sequence signature, consistent with CLEs arising repeatedly from otherwise typical intronic sequence rather than representing a conserved sequence class.

Together, these analyses show that CLE sequences are poorly conserved, show limited cross-species alignment, and retain the TE characteristics of their host introns, distinguishing them from conventional alternative last exons and indicating that most CLEs are predominantly species-specific products of recent exonisation.

### <u>CLE inclusion is developmentally regulated and inversely associated with</u> <u>full-length splice usage.</u>

If CLE inclusion represents a regulated component of transcript processing rather than stochastic transcriptional noise, its usage should vary reproducibly during development and be coupled to the competing full-length (FL) splice outcome. To test this, we quantified CLE and FL splicing across developmental RNA-seq series from zebrafish, mouse, and human. All three publicly available datasets were generated using poly(A)-selected libraries (Supplementary Datasheet 6)^31,32^, allowing CLE- and FL-containing transcripts to be assessed using equivalent sequencing methods (Figure 3A).

**Figure 3:**
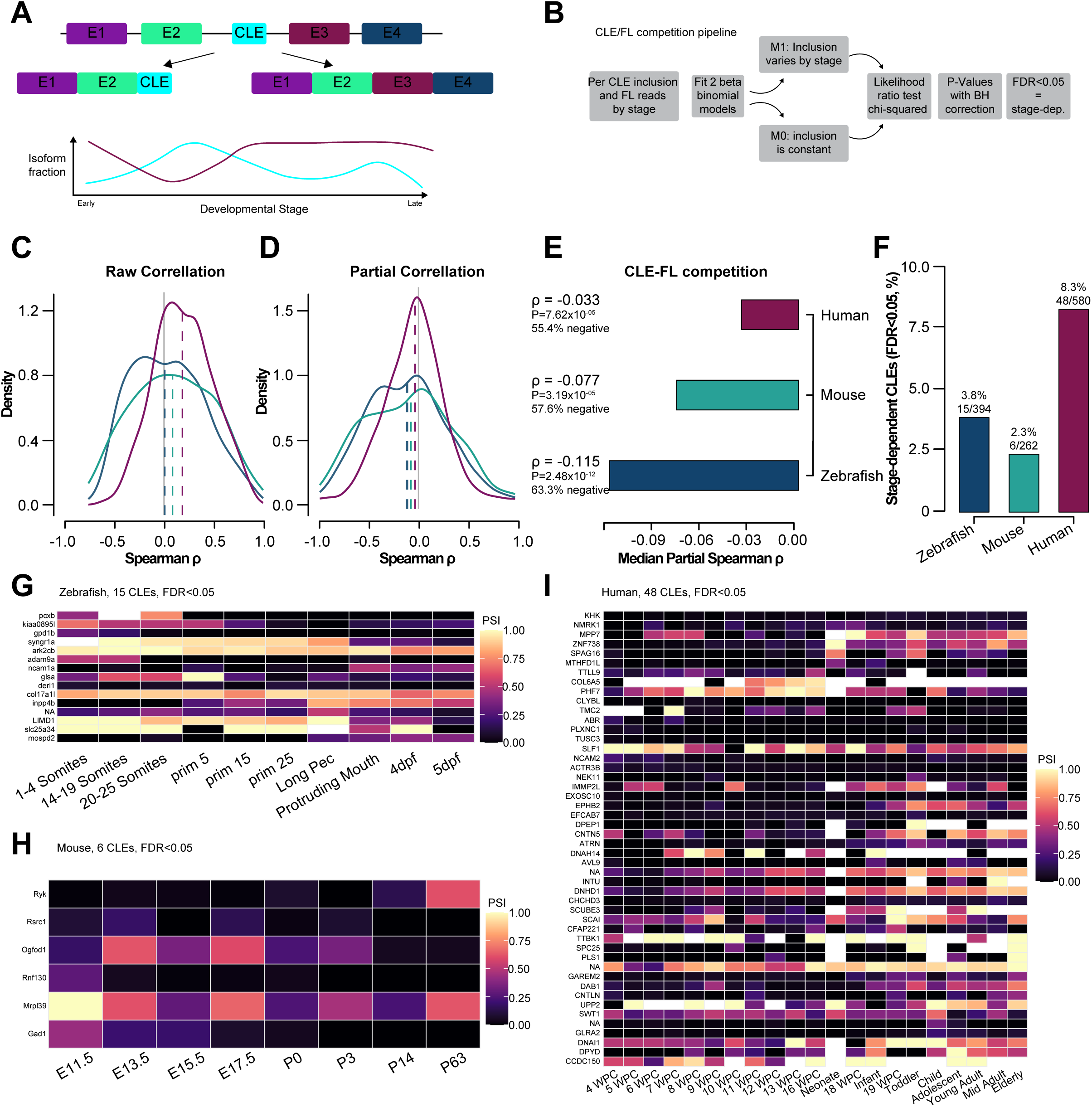
CLE inclusion is developmentally regulated and inversely associated with competing terminal-splice usage. **(A)** Schematic illustrating competition between CLE inclusion and continued downstream splicing from a shared upstream donor during development. **(B)** Analysis pipeline used to identify developmentally regulated CLEs. Replicate-level CLE and competing splice-junction counts were modelled using beta-binomial models in which CLE inclusion was either allowed to vary with developmental stage or constrained to remain constant. Models were compared by likelihood-ratio test and P values were corrected using the Benjamini–Hochberg method. **(C)** Distribution of raw Spearman correlations between CLE and competing non-CLE splice-junction usage across developmental stages in zebrafish, mouse, and human. Dashed lines indicate species medians. **(D)** Distribution of partial Spearman correlations between CLE and competing splice usage after conditioning on upstream-exon read count as a proxy for host-gene RNA abundance. **(E)** Median partial Spearman correlations were negative in all three species: zebrafish, ρ=−0.115, 63.3% negative, P=2.48×10⁻¹²; mouse, ρ=−0.077, 57.6% negative, P=3.19×10⁻⁵; human, ρ=−0.033, 55.4% negative, P=7.62×10⁻⁵; two-sided one-sample Wilcoxon signed-rank tests. **(F)** Percentage and number of testable CLEs showing significant developmental-stage dependence at FDR<0.05: zebrafish, 15/394 (3.8%); mouse, 6/262 (2.3%); human, 48/580 (8.3%). **(G–I)** Stage-resolved CLE PSI for CLEs showing significant developmental regulation in zebrafish (G), mouse (H), and human (I). Black cells indicate stages in which PSI was not measurable.

CLE inclusion and FL splicing were quantified directly from splice-junction reads. Because a CLE and its competing FL splice isoform originate from the same upstream donor, reads spanning this donor were classified according to whether they terminated at the CLE acceptor or at a downstream non-cryptic acceptor. Rather than defining the competing FL splice solely from the reference annotation, we identified the dominant non-CLE acceptor used from each donor empirically across the corresponding RNA-seq dataset. Each CLE was therefore compared with the full-length splice outcome that most frequently competed for the same donor in the tissues analysed. In parallel, read coverage across the upstream exon was quantified as a proxy for host-gene transcription that does not depend on the downstream splice choice (Figure 3B).

We first examined the relationship between CLE and FL splice usage across development. For each CLE, junction counts were averaged across biological replicates within each developmental stage, and the relationship between CLE and FL usage was then assessed across stages. Before accounting for host-gene transcription, CLE and FL junction counts showed weak positive correlations, with median Spearman correlations of +0.007 in zebrafish, +0.082 in mouse, and +0.184 in human (Figure 3C, Supplementary Datasheet 6). This positive relationship is expected if both splice products vary with the overall transcriptional output of their shared host gene. We therefore calculated partial Spearman correlations between CLE and FL splice usage while conditioning on upstream-exon coverage. This asks whether the two alternative splice outcomes remain associated after accounting statistically for developmental variation in host-gene transcription.

After accounting for this shared transcriptional variation, CLE and FL splice usage showed a consistent negative relationship in all three species (median partial Spearman ρ = −0.115 in zebrafish, −0.077 in mouse, and −0.034 in human; Figure 3D-E, Supplementary Datasheet 6). Most CLEs showed a negative transcription-corrected relationship with their competing FL splice isoform (63.3% in zebrafish, 57.6% in mouse, and 55.4% in human, Figure 3D-E), and the partial-correlation distributions were significantly shifted below zero in each species (Wilcoxon signed-rank test: zebrafish, P = 2.5×10⁻¹², n = 455; mouse, P = 3.2×10⁻⁵, n = 278; human, P = 7.6×10⁻⁵, n = 601, Figure 3E, Supplementary Datasheet 6). Thus, after accounting for variation in host-gene transcription, increased CLE usage is associated with reduced usage of the competing FL splice isoform. This inverse relationship is consistent with competition between alternative terminal-splicing outcomes from a shared donor.

We next asked directly whether CLE inclusion changes across developmental time. For each testable CLE, stage-resolved CLE and FL junction counts from individual biological replicates were modelled using a beta-binomial framework that accommodates overdispersion between replicates (Figure 3B). We compared a model in which CLE inclusion was allowed to vary with developmental stage against a null model in which inclusion remained constant. Likelihood-ratio P values were corrected across all tested CLEs using the Benjamini-Hochberg procedure.

A significant developmental-stage effect was detected for a defined subset of CLEs in every species examined: 15 of 394 CLEs in zebrafish (3.8%), 6 of 262 in mouse (2.3%), and 48 of 580 in human (8.3%) at FDR < 0.05 (Figure 3F, Supplementary Datasheet 6). Individual stage-regulated CLEs displayed distinct developmental trajectories, including transient, early-enriched, and later-enriched patterns of inclusion (Figure 3G-I, Supplementary Datasheet 6). Thus, CLE usage is not universally dynamic, but a reproducible subset of CLE splice choices shows significant stage dependence across development in each vertebrate dataset.

Together, these analyses identify two complementary properties of physiological CLE usage. First, across the broader CLE population, CLE usage is inversely associated with use of the competing full-length splice after accounting for variation in host-gene transcription. Second, individual CLEs can undergo statistically significant changes in splice-site usage across development. Developmental stage dependence and an inverse relationship with full-length splice usage are therefore recurrent features of CLE processing across vertebrate development, consistent with regulated competition between alternative terminal-splicing outcomes.

### <u>CLEs previously revealed by SFPQ loss are expressed during normal</u> <u>zebrafish development</u>

Having established developmental regulation of CLE inclusion quantitatively using RNA-seq, we next asked whether CLEs originally identified following loss of the splicing regulator SFPQ are also used during normal development. Previous work identified 113 CLEs in *sfpq*^-/-^ zebrafish. We therefore performed an orthogonal RT-PCR screen using RNA collected from wild-type zebrafish at 1, 2, 3, 4, and 5dpf, with primers spanning the upstream exon-to-CLE splice junction (Supplementary Figure 4A). In parallel, we generated deep RNA-seq from wild-type zebrafish at 1dpf and quantified reads spanning each CLE splice junction, providing an independent sequencing-based readout of CLE production during normal development.

Of the 113 previously identified SFPQ-sensitive CLEs, 83 yielded a correctly sized RT-PCR product from *sfpq*^-/-^ cDNA and were therefore considered technically validated for the screen. Of these 83 primer-validated CLEs, 74 (89.2%) were detected at one or more developmental stages in wild-type embryos or larvae, whereas nine were not detected under wild-type assay conditions (Supplementary Figures 3A and 4A-B). The remaining 30 assays did not generate the expected product from *sfpq^-/-^* positive-control cDNA and were therefore considered technically uninformative rather than classified as absent from wild-type animals. Thus, among CLEs for which primer performance could be independently confirmed, the large majority were detectable during normal zebrafish development.

**Figure 4:**
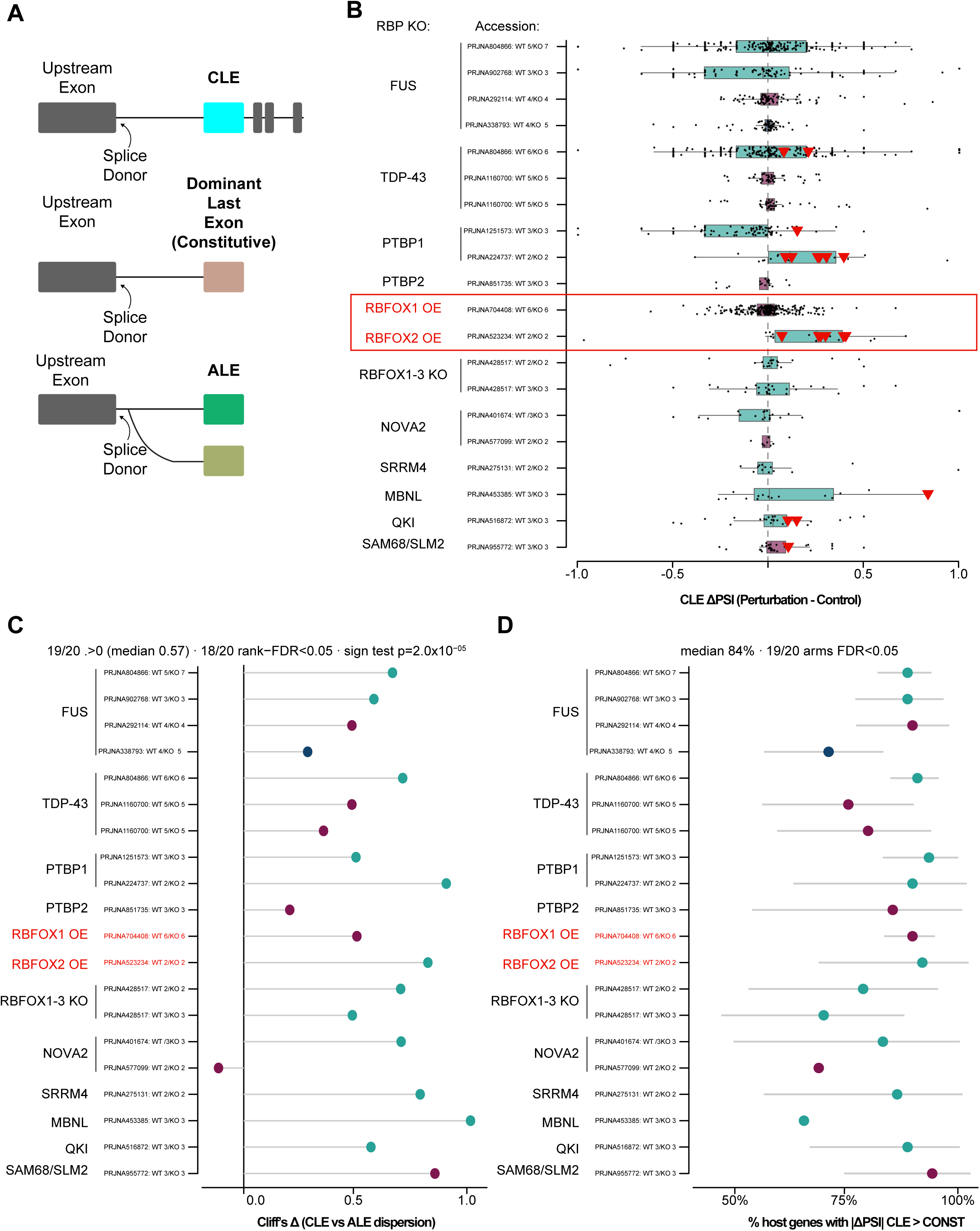
CLE usage is preferentially labile following perturbation of neural RNA-binding proteins. **(A)** Schematic of the three terminal-exon classes compared across RBP perturbation datasets: CLEs, dominant annotated last exons (CONST), and conventional alternative last exons (ALEs). Event usage was quantified from splice-junction reads arising from each event’s upstream donor. **(B)** Distribution of CLE ΔPSI values, calculated as mean PSI in the perturbed condition minus mean PSI in the corresponding control, across 20 RBP perturbation arms spanning zebrafish, mouse, and human. Points represent individual CLEs and box plots show the distribution within each perturbation arm. Red triangles indicate 19 CLEs classified as emergent from an effectively silent state in the control condition. **(C)** Effect size for CLE-versus-ALE perturbation-associated dispersion after one-to-one matching for baseline control PSI and donor-junction depth. Cliff’s δ was positive in 19/20 arms (median δ=0.57), with significantly greater CLE dispersion in 18/20 arms after Benjamini–Hochberg correction using the median-centred rank-based dispersion test (across-arm sign test, P=2.0×10⁻⁵). **(D)** Percentage of CLE-host genes in which the absolute CLE perturbation response exceeded that of the dominant annotated last exon from the same gene. The primary analysis was restricted to genes containing a single testable CLE. Across the panel, the median proportion was 84%, with paired CLE-versus-CONST differences significant in 19/20 perturbation arms after Benjamini–Hochberg correction.

Independent analysis of the 1 dpf RNA-seq data provided a second measure of physiological CLE production by quantifying reads spanning the upstream exon-to-CLE splice junction (Supplementary Datasheet 7). This sequencing-based analysis recovered CLE junctions for 63 of the of SFPQ-sensitive events at this single developmental stage, including 20 events not detected by RT-PCR under the corresponding assay conditions (Supplementary Datasheet 7). The incomplete overlap between the two approaches is consistent with their different detection characteristics and, in the case of RT-PCR, the broader developmental sampling window. Considered together, these orthogonal approaches demonstrate that CLEs originally revealed following loss of SFPQ are frequently produced in the wild-type transcriptome. The RT-PCR and 24 hpf RNA-seq screens were therefore used as qualitative evidence of physiological CLE production, whereas quantitative evidence for developmental regulation is provided by the stage-resolved RNA-seq analysis above.

The most abundant CLE previously identified following SFPQ loss arises from the *ephA4b* gene. *ephA4b-CLE* was readily detected by RT-PCR throughout the first five days of wild-type development (Supplementary Figure 4) and reads spanning the upstream exon-to-CLE splice junction were consistently recovered across replicate wild-type 24 hpf RNA-seq libraries (Supplementary Datasheet 7), independently demonstrating production of the spliced CLE transcript. We further examined the anatomical distribution of the CLE-containing region using hybridisation chain reaction (HCR). CLE-region signal was detected within the broader anatomical expression domain of *ephA4b*, including developing neural structures (Supplementary Figure 3B). To distinguish mature *ephA4b-CLE* RNA from nascent pre-mRNA, CLE-targeting HCR probes were used together with a second probe set targeting the downstream sequence of the host intron (Supplementary Figure 3C–E). Puncta detected by the CLE probes but lacking coincident intronic signal were therefore classified as CLE-specific rather than intron-containing pre-mRNA. In single optical sections, some CLE-positive puncta overlapped DAPI-positive nuclei (white circles), consistent with sites of transcription or nuclear RNA, whereas others were clearly spatially separated from DAPI-positive nuclear regions (red circles; Supplementary Figure 3D). The presence of these non-intronic, extranuclear CLE puncta provides spatial evidence for mature *ephA4b-CLE* transcripts in wild-type embryos, complementing the independent splice-junction RT-PCR and RNA-sequencing evidence for the mature isoform.

CLE-specific puncta were identified at 24, 36 and 48 hpf using the same non-overlap criterion (Supplementary Figure 3E), supporting physiological production of *ephA4b-CLE* during the period of retinal and neural development in which its function was subsequently examined. Together, these observations show that *ephA4b-CLE* production is not restricted to SFPQ-deficient conditions but occurs during normal zebrafish development.

Overall, CLEs originally exposed by loss of SFPQ are frequently detectable in wild-type animals using independent molecular approaches. These observations support a model in which SFPQ suppresses or modulates splice events that can also form part of the physiological transcript repertoire, rather than generating an entirely novel class of transcripts only after splicing-factor dysfunction.

### <u>Neural RNA-binding-protein perturbation reveals preferential lability of</u> <u>CLE usage.</u>

Having established that CLE inclusion changes dynamically during normal vertebrate development, we next asked whether CLE usage is sensitive to changes in the trans-acting splicing environment We assembled a panel of 20 independent, neural RBP perturbation comparisons comprising loss-of-function or depletion experiments, RBP overexpression, and disease-associated RBP variants across mouse, human and zebrafish, including FUS, TDP-43, PTBP1/2, RBFOX1/2, NOVA2, SRRM4, MBNL, QKI and SAM68/SLM2 (Figure 4A-B, Supplementary Datasheet 8). Where the same RBP appears in more than one comparison, these represent distinct experimental datasets, perturbation contexts or conditions and were analysed independently. Because perturbation of different RBPs, and distinct perturbations of the same RBP, can alter individual splice targets in opposite directions, our subsequent analysis tested the magnitude and dispersion of CLE responses rather than requiring a common direction of change. Within each experiment, CLEs were compared with two classes of annotated terminal-exons: the constitutive terminal exon (CONST; operationally defined as the dominant annotated last exon) and conventional alternative last exons (ALEs). For each event, usage was quantified directly from splice-junction reads arising from the shared upstream splice donor, i.e. the 5′ splice site at the end of the exon immediately upstream of the terminal exon (Figure 4A, Supplementary Datasheet 8), and the change in usage (ΔPSI) was defined as mean donor-specific PSI in the perturbed condition minus that in the corresponding control.

Across RBP perturbation comparisons, CLE ΔPSI responses were heterogeneous in both magnitude and direction (Figure 4B, Supplementary Datasheet 8). Interestingly, individual RBP perturbations produced both increased and decreased CLE usage (Figure 4B, Supplementary Datasheet 8). Overall, the RBP perturbation panel showed no uniform direction of effect with the exception of RBFOX2 overexpression which produced a predominantly positive shift and PTBP1 depletion which produced a predominantly negative shift in CLE usage (Figure 4B, Supplementary Datasheet 8). Perturbation of neural RBP state therefore does not simply increase or decrease CLE inclusion. Instead, CLE splice choice responds in an RBP- and target-dependent manner.

We next investigated whether the change in CLE usage (ΔPSI) differed from that of control exon type ALEs and dominant last exons (constitutive exons). Because opposing responses can cancel in analyses of mean ΔPSI, we next tested whether RBP perturbation altered the dispersion of CLE usage relative to ALEs, thereby quantifying perturbation-associated lability irrespective of direction. To determine whether this lability was specific to CLE usage rather than a general property of alternative terminal splicing, we compared CLEs with conventional ALEs. Within each perturbation comparison, CLE and ALE events were matched one-to-one for both donor-junction coverage and baseline control PSI using nearest-neighbour matching without replacement (Figure 4C, Supplementary Datasheet 8). Matching baseline PSI was important because it constrained the available dynamic range of the two event classes, while matching donor depth reduced the possibility that greater CLE variability simply reflected lower junction coverage. We then compared the absolute deviations of CLE and ALE ΔPSI values from their respective class medians using a median-centred rank-based dispersion test. CLE usage was significantly more dispersed than that of baseline- and depth-matched ALEs in 18 of 20 perturbation comparison after Benjamini–Hochberg correction (FDR < 0.05, Figure 4C, Supplementary Datasheet 8). The excess CLE dispersion was highly consistent across the panel: Cliff’s δ favoured greater CLE dispersion in 19 of 20 comparisons, with a median δ of 0.57 (across-comparison sign test, P = 2.0 × 10⁻⁵), and every comparison showed a CLE-to-ALE variance ratio greater than one, with a median ratio of approximately 3.7-fold (Figure 4C, Supplementary Datasheet 8). Independent Brown–Forsythe and Fligner–Killeen dispersion tests showed the same pattern of greater CLE dispersion and provided conservative sensitivity analyses (Supplementary Datasheet 8).

Because ALEs are themselves regulated alternative terminal exons, this comparison shows that CLE usage is unusually labile even relative to conventional alternative terminal-splice choices. Importantly, this conclusion does not depend on comparing low-usage CLEs with near-saturated constitutive exons. This CLE vs ALE analysis explicitly controls for both baseline usage and donor-junction coverage. Thus, increased CLE responsiveness cannot readily be attributed to differences in read depth or to the greater mathematical range available to low-PSI events.

We next asked whether the excess CLE response could instead arise because CLEs preferentially occur in genes that are themselves unusually sensitive to RBP perturbation. To control directly for host-gene identity, we compared each CLE with the constitutive terminal exon from the same gene (Figure 4D). We restricted the test to genes containing a single testable CLE and found that the absolute perturbation-associated change in CLE usage exceeded that of the gene’s own constitutive terminal exon in a median of 84% of host genes across the panel, with individual-comparison proportions ranging from 64% to 90% (Figure 4D, Supplementary Datasheet 8). Paired differences were significant in 19 of 20 perturbation comparisons after FDR correction (paired Wilcoxon signed-rank test, Figure 4D, Supplementary Datasheet 8).

Together, the ALE and matched constitutive last exon comparisons suggest that the CLE inclusion effect is not explained by lower CLE junction coverage, differences in baseline PSI, or by preferential localisation of CLEs in unusually RBP-responsive genes. Instead, CLE usage shows a greater perturbation-associated lability than both ALEs and constitutive last exons.

Finally, we applied the same emergence criterion across all 20 RBP perturbation comparisons, irrespective of perturbation class, to identify CLEs that transitioned from an effectively silent state under control conditions to reproducible inclusion following perturbation. Emergent CLEs were identified following both loss-of-function/depletion and overexpression perturbations (red triangles, Figure 4B; Supplementary Datasheet 8). Emergence was defined independently of event-level significance testing, and an event was required to have informative donor coverage in at least two biological replicates per condition, PSI < 0.02 in every informative control replicate, and PSI ≥ 0.05 with direct target-junction support in a strict majority of informative perturbed replicates. Using this replicate-defined criterion, we identified 20 clean emergent CLEs across seven perturbation comparisons. These included *Dpp10* following MBNL depletion, which showed the largest observed transition (ΔPSI ≈ 0.84), as well as events in *Large1* and *Twsg1* following RBFOX2 overexpression, *Epha10* and *Mtcl3* following PTBP2 depletion, and *Camk2g* following TDP-43 LoF (Supplementary Datasheet 8). These events demonstrate that altered RBP state can expose CLE choices that are effectively silent under the corresponding control condition.

Together, these analyses identify CLE inclusion as an unusually labile terminal-splicing outcome. Across neural RBP perturbations, CLE usage showed larger and more dispersed changes than baseline- and coverage-matched conventional ALEs, and this excess response persisted when CLEs were compared with the constitutive terminal exon within the same host genes. A subset of CLEs additionally transitioned from an effectively silent state to reproducible inclusion following perturbation. Cryptic terminal-exon choice therefore represents a dynamically RBP-sensitive component of terminal-exon regulation rather than a fixed, low-level by-product of transcription.

### A subset of CLEs is translated during normal vertebrate development

Since CLEs show specific temporal expression patterns, exert a subtle effect on full-length transcription, and are regulated by RBPs, we next asked whether CLEs are also translated. The detection of many CLE mRNAs across zebrafish, mouse, and human suggests a functional role, exerted either through altering full-length transcription or through translation into CLE proteins. Indeed, in *sfpq^-/-^* zebrafish mutants, the deleterious effects of the ephA4b-CLE are abolished when its translation is blocked, indicating that its function is translation-dependent^19^. We therefore asked whether CLE-specific peptides could be detected more broadly in publicly available mass spectrometry datasets (Supplementary Datasheet 9). Non-canonical translation has been surveyed previously^33^, but these studies did not examine CLEs, and because CLEs are predominantly low-inclusion isoforms, untargeted searches would be unlikely to recover them.

To determine whether any CLEs are translated, we mined publicly available label-free data-dependent acquisition (DDA) mass-spectrometry datasets derived from CNS tissues across zebrafish, mouse and human (Supplementary Datasheet 9). If CLE reading frames are translated, peptides unique to them should be recoverable from these data. We searched a panel of 84 deep CNS proteomes, spanning wild-type/developmental and disease/disease-model contexts, for CLE-specific peptides (Figure 5A and Supplementary Datasheet 9). To control for the low complexity of some cryptic sequences, we applied two independent detection strategies: a conventional target-decoy search with Percolator 1% FDR control (Supplementary Figure 5A), and a stringent composition-matched decoy-margin test in which each peptide was required to outscore eight length- and composition-matched decoys and to exceed the highest score reached by any decoy in its species.

**Figure 5:**
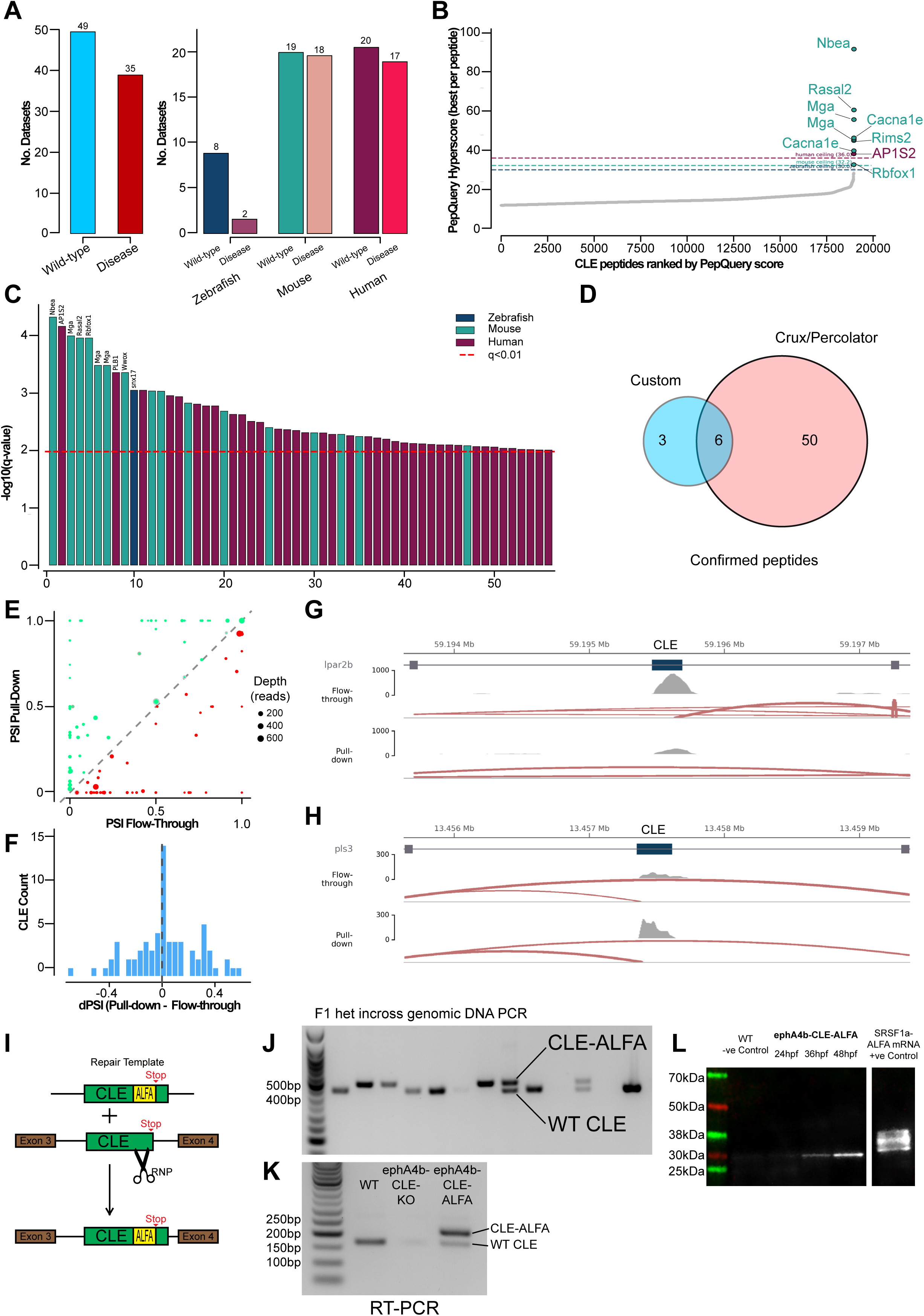
A subset of CLE-containing transcripts associates with ribosomes and can generate stable protein products *in vivo*. **(A)** Public CNS proteomic datasets analysed across zebrafish, mouse, and human, separated into wild-type/developmental and disease-associated datasets. **(B)** CLE-specific peptides ranked according to their maximum PepQuery hyperscore. Dashed lines indicate the species-specific maximum score observed among eight composition-matched decoy peptide sets. Labelled peptides exceeded the corresponding species-wide decoy-score ceiling. **(C)** CLE-specific peptides identified by Crux/Percolator ranked by peptide-level significance. Dashed line indicates q=0.01; colours indicate species. **(D)** Overlap of CLE peptide detections obtained using the composition-matched PepQuery strategy and the Crux/Percolator search. **(E)** CLE isoform partitioning between TRAP fractions. CLE inclusion (PSI) in the ribosome pull-down versus flow-through is shown for each testable CLE; points above the diagonal indicate higher CLE inclusion in the pull-down, and point colour indicates ΔPSI (pull-down − flow-through). **(F)** Distribution of ΔPSI (pull-down − flow-through) across CLEs; of 130 testable CLEs, 51 showed higher inclusion in the pull-down (ΔPSI > 0) and 34 in the flow-through (ΔPSI < 0). **(G–H)** Genome-browser coverage and splice-junction tracks for lpar2b-CLE (G), preferentially retained in the flow-through, and pls3-CLE (H), preferentially included in the ribosome pull-down, shown for representative flow-through and pull-down samples. **(I)** Schematic of the CRISPR/Cas9-mediated endogenous *ephA4b-CLE* ALFA-tagging strategy, in which the ALFA epitope was inserted immediately upstream of the CLE-encoded stop codon. **(J)** PCR genotyping of genomic DNA from an F1 *ephA4b-CLE-ALFA* heterozygous in-cross. Wild-type and ALFA knock-in alleles produce 436-bp and 475-bp products, respectively. **(K)** RT-PCR confirming expression of the *ephA4b-CLE-ALFA* transcript. **(L)** Anti-ALFA western blot of wild-type and homozygous *ephA4b-CLE-ALFA* embryo lysates at 24, 36, and 48 hpf. A band at the predicted truncated EphA4b-CLE size (∼31–32 kDa) was detected at 36 and 48 hpf and was absent from wild-type controls. SRSF1a-ALFA mRNA-injected embryos were used as a positive control.

As expected for a predominantly low-usage splice-isoform class measured in complex tissues, few CLE-specific peptides were confidently detected. This limited recovery is not unexpected in data-dependent acquisition (DDA) proteomics, in which precursor selection favours relatively abundant peptides: CLE-specific peptides derived from low-abundance or cell-type-restricted isoforms may therefore be infrequently sampled for fragmentation. Of 18,957 CLE peptides evaluated, 304 produced PepQuery matches; however, most scored no higher than their composition-matched decoys, whereas peptides from the corresponding full-length isoforms were readily recovered. Thus, failure to detect a CLE-specific peptide cannot distinguish absence of translation from abundance below the sensitivity of shotgun proteomics, including as a consequence of cell-type-restricted expression (Figure 5B and Supplementary Figure 5A). The decoy margin test identified nine CLE peptides, from seven genes (Nbea, Rasal2, Mga, Cacna1e, Rims2, AP1S2, and Rbfox1), that scored above the species-wide decoy ceiling (Figure 5B). The conventional Crux/Percolator search recovered 56 cryptic peptides at 1% FDR (Figure 5C and Supplementary Datasheet 9). Five of the seven above-ceiling genes, Nbea-CLE, Rasal2-CLE, Mga-CLE, AP1S2-CLE and Rbfox1-CLE (the last itself a neuronal splicing regulator), were also recovered by the conventional search, representing the highest-confidence, cross-method-confirmed detections (Figure 5D); the remaining two, Cacna1e-CLE and Rims2-CLE, were supported by the decoy-margin test alone. The Nbea-CLE peptide was the strongest, scoring above its decoys in three independent mouse datasets (Supplementary Datasheet 9). Because the datasets used span different tissues, developmental stages, and conditions, each CLE is expected to appear only in the subset of datasets matching its expression; detection in even a single dataset above the decoy null is therefore meaningful, and recurrence across datasets, as observed for Nbea-CLE, further strengthens the result. The confirmed peptides were detected across both wild-type/developmental and disease-associated proteomes, with the highest-confidence detections found predominantly in normal and developmental datasets (Supplementary Figure C), consistent with constitutive translation of these CLE isoforms during normal development rather than a disease-specific phenomenon. Public proteomes therefore confirm CLE translation for a defined set of genes but are limited by depth for the low-abundance majority.

To assess whether physiological CLE-containing transcripts associate with the translational machinery, we performed Translating Ribosome Affinity Purification (TRAP), a method previously used in zebrafish to isolate ribosome-associated mRNAs^26^. We injected mRNA encoding *rpl10a-3xHA-P2A-EGFP* into zebrafish embryos and isolated anti-HA TRAP pull-down and corresponding flow-through RNA at 1, 2 and 3 dpf in three biological replicates (Supplementary Figure 5D-H). In total, 162 CLE-containing transcripts were detected across the two fractions (Supplementary Datasheet 10). Because whole-embryo poly(A) TRAP saturates the pull-down with the abundant embryonic translated pool, we quantified CLE association at the isoform level as the change in CLE inclusion between fractions (ΔPSI = pull-down − flow-through). Of 130 CLEs testable at the splice-junction level, 51 showed higher CLE inclusion in the ribosome pull-down and 34 in the flow-through (Figure 5E–F, Supplementary Datasheet 10), identifying subsets of CLE-containing transcripts with preferential ribosomal or non-ribosomal association. Representative examples are shown for *lpar2b-CLE*, which is preferentially retained in the flow-through (Figure 5G), and *pls3-CLE*, which is preferentially included in the ribosome pull-down (Figure 5H). Consistent with ribosome-selective capture, non-coding lncRNAs were depleted from the pull-down while CLE-host genes partitioned with translated protein-coding mRNA (Wilcoxon P = 2×10⁻⁴; Supplementary Figure 5I). These data identify a subset of physiological CLE-containing transcripts that preferentially associate with ribosomes during zebrafish development.

Ribosome association does not itself demonstrate completion of translation or production of a stable protein. We therefore sought direct evidence for translation of an endogenous CLE-containing transcript. We focused on *ephA4b-CLE* because it is strongly induced following loss of SFPQ^19^ and is independently detectable during normal zebrafish development by splice-junction RT-PCR and RNA-seq. This provided a well-characterised endogenous locus at which to test directly whether a physiological CLE transcript generates a protein.

We generated an endogenous *ephA4b-CLE* knock-in line in which an ALFA epitope^34^ was inserted immediately upstream of the CLE-encoded stop codon by CRISPR/Cas9-mediated homology-directed repair using a donor carrying 40-bp homology arms (Figure 5H). Genomic insertion was validated by PCR and sequencing of F1 animals (Figure 5I), and production of the *ephA4b-CLE-ALFA* transcript was confirmed by RT-PCR (Figure 5J). Western blotting of homozygous *ephA4b-CLE-ALFA* embryo lysates revealed a band at the predicted truncated protein size of approximately 31–32 kDa at 36 and 48 hpf that was absent from wild-type controls (Figure 5K). This provides direct evidence that an endogenous CLE-containing transcript can produce a stable truncated protein during normal development. Together, the TRAP analysis, public proteomic detections and endogenous *ephA4b-CLE* tagging support translational engagement of a subset of CLE transcripts, while demonstrating directly that at least one physiological CLE isoform produces a stable protein *in vivo*.

### <u>Convergent EphA4 CLEs generate similar predicted ligand-binding</u> <u>ectodomains</u>

The evolutionary analyses above suggested that orthologous genes can repeatedly acquire CLEs despite little conservation of the underlying cryptic sequence. We therefore asked whether independently arising CLEs could nevertheless converge on similar protein-level consequences. *EphA4* provided a striking example. We identified two CLEs in human *EPHA4* and one CLE each in the zebrafish paralogues *ephA4a* and *ephA4b*. In each case, the CLE occurs within the orthologous intron immediately downstream of the same conserved coding exon, whereas no equivalent CLE was detected in mouse *Epha4* despite preservation of the surrounding exon architecture (Supplementary Figure 6A). Despite arising from different cryptic sequences, the resulting transcripts terminate the EphA4 coding sequence at a similar position within the extracellular region. The predicted proteins retain the signal peptide, complete ephrin-binding domain and part of the cysteine-rich region, while removing both fibronectin type III repeats, the transmembrane segment and the intracellular kinase, SAM and PDZ-binding regions (Supplementary Figure 6B). Thus, distinct EphA4 CLEs converge on a similar predicted ectodomain architecture.

We next tested whether CLE-mediated truncation was predicted to disrupt folding of the retained receptor region or its interaction with ephrin ligands. Full-length and CLE-derived EphA4 proteins were modelled in complex with species-matched ephrin-A2 using AlphaFold3 (Supplementary Figure 6C–E; Supplementary Datasheet 11). In each species, the retained CLE-derived receptor region closely superimposed on the corresponding region of the full-length receptor, with backbone RMSDs of 0.54 Å for human EPHA4, 0.60 Å for zebrafish EphA4a and 0.87 Å for zebrafish EphA4b (Supplementary Figure 6G). Isolated CLE-derived proteins were also predicted to fold confidently as monomers, with pTM values of 0.71–0.81.

The predicted ephrin interaction was similarly preserved following CLE-mediated truncation. Interface-confidence scores were comparable between CLE and full-length complexes, with ipTM values of 0.66 versus 0.68 for human EPHA4, 0.69 versus 0.69 for zebrafish EphA4a and 0.75 versus 0.69 for zebrafish EphA4b (Supplementary Figure 6F). CLE complexes also showed lower predicted interface PAE than the corresponding full-length models (human, 1.13 versus 1.50 Å; EphA4a, 1.19 versus 1.76 Å; EphA4b, 1.09 versus 1.64 Å; Supplementary Figure 6H). Receptor residues predicted to contact ephrin were confined to the retained ligand-binding domain and were preserved in the CLE-derived structures (Supplementary Figure 6I; Supplementary Datasheet 11). Together, these analyses show that independently arising EphA4 CLEs converge on proteins predicted to retain a folded ephrin-binding ectodomain while lacking the membrane anchor and intracellular signalling machinery of the full-length receptor. This architecture is compatible with a soluble, secreted mode of action, although secretion, ligand affinity and competitive antagonism remain to be established experimentally.

### <u>Expression of *ephA4b-CLE* is required for normal development of the</u> <u>visual system.</u>

If CLE mRNAs play functional roles during development, loss of a specific CLE should result in a defined developmental defect. To investigate whether CLEs have functional roles, we assessed whether *ephA4b-CLE* was needed for normal visual development, given its dynamic expression in the developing retinas and ventral diencephalon. EphA4 signalling regulates optic nerve growth by controlling retinal ganglion cell (RGC) axon guidance and midline crossing^35–37^. We therefore examined whether loss of *ephA4b-CLE* expression affects RGC development, following RGC differentiation and the optic nerve over time using *tg(ath5:RFP)* transgenic background (expressing RFP in RGCs and RGC-derived optic nerve).

We generated a CRISPR/Cas9 targeted deletion of the *ephA4b* intronic *CLE* sequence using two ribonucleoprotein (RNP) complexes whose cut-sites flanked the CLE (Figure 6A). Deletions in founders were validated by PCR using genomic DNA and primers flanking the cut sites (436bp wild-type band, 121bp mutant band, Figure 6B). Mutant alleles were confirmed by sequencing and established F1 heterozygotes were crossed to generate homozygous *ephA4b^ΔCLE/ΔCLE^,* heterozygous *ephA4b^ΔCLE/+^,* and wild-type *ephA4b ^+/+^* sibling control populations. We validated loss of *ephA4b-CLE* expression using RT-qPCR and showed ∼50% reduction of *ephA4b-CLE* mRNA abundance in *ephA4b^ΔCLE/+^* fish and was not detectable in *ephA4b^ΔCLE/ΔCLE^* fish (N=6, 50 embryos for each extraction, P<0.0001, Ordinary one-way ANOVA with Tukey’s multiple comparisons; Figure 6C). We also showed that expression of the full-length transcript for the *ephA4b* gene is unaffected in *ephA4b^ΔCLE/+^* and *ephA4b^ΔCLE/ΔCLE^* fish (N=6, 50 embryos for each extraction, not significant, Ordinary one-way ANOVA with Tukey’s multiple comparisons; Figure 6D).

**Figure 6:**
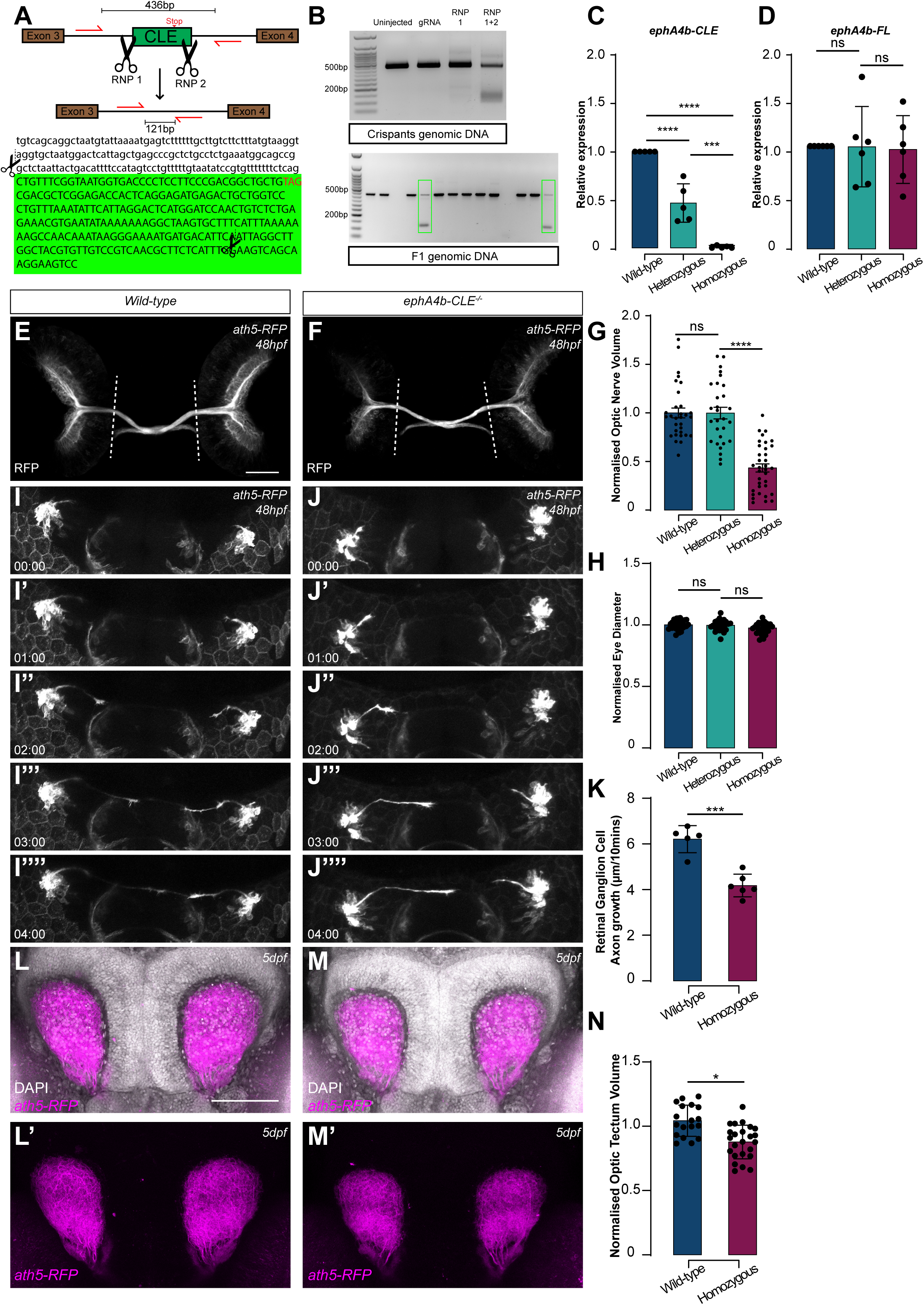
Loss of ephA4b-CLE impairs retinal ganglion cell axon growth and projection. **(A)** Schematic of the CRISPR/Cas9 strategy used to generate the stable *ephA4b*^ΔCLE^ allele. Two RNP complexes flanking the CLE removed a 315-bp region of *ephA4b* intron 3 containing the CLE; wild-type and deletion alleles generate 436-bp and 121-bp PCR products, respectively. **(B)** PCR validation of the CLE deletion in F0 crispants and germline-transmitting F1 animals. **(C–D)** RT-qPCR measurement of *ephA4b-CLE* (C) and full-length *ephA4b* (D) transcript abundance in 24-hpf wild-type, *ephA4b^ΔCLE/+^* and *ephA4b^ΔCLE/ΔCLE^*embryos. *ephA4b-CLE* abundance was reduced in heterozygotes and undetectable in homozygotes, whereas the assayed full-length transcript was unchanged (N=6 biological replicates per genotype; 50 embryos per replicate; ordinary one-way ANOVA with Tukey’s multiple comparisons). **(E–F)** Maximum-intensity projections of 48-hpf wild-type (E) and *ephA4b^ΔCLE/ΔCLE^*(F) *tg(ath5:RFP)* embryos showing RGC axons and optic nerves. Dashed lines indicate the region quantified. **(G)** Normalised optic-nerve volume in wild-type (N=31), heterozygous (N=29), and homozygous (N=35) embryos. Optic-nerve volume was reduced in *ephA4b^ΔCLE/ΔCLE^* animals; linear mixed-effects model, ****P<0.0001. **(H)** Normalised eye diameter in wild-type (N=30), heterozygous (N=24), and homozygous (N=28) embryos; no significant differences were detected. **(I–J)** Time-lapse confocal imaging of pioneer RGC axon growth in wild-type (I–I’’’’) and *ephA4b^ΔCLE/ΔCLE^*(J–J’’’’) embryos over 4 h. **(K)** Mean pioneer RGC axon growth per 10-min interval in wild-type and homozygous mutants. **(L–M)** Representative dorsal views of RGC projections into the optic tectum at 5 dpf in wild-type (L–L′) and *ephA4b^ΔCLE/ΔCLE^* (M–M′) larvae; DAPI is shown in grey and ath5:RFP-positive projections in magenta. **(N)** Normalised optic-tectum projection volume at 5 dpf was reduced in homozygous mutants; Welch’s t-test, *P<0.05. Data are shown as mean ± SD. Scale bars = 50µm.

At 48hpf, the RGC axon projections were visualised in fixed embryos by confocal microscopy and total volume of the optic nerves that had exited the retinas (between dashed lines, Figure 6E, F) was quantified. In *ephA4b^ΔCLE/ΔCLE^* embryos, the optic nerve volume was reduced compared to the wild-type volume (1.000 normalised volume in wild-type; N=31, 0.998 in *ephA4b^ΔCLE/+^*; N=29, and 0.433 in *ephA4b^ΔCLE/ΔCLE^*; N=35, P<0.0001, linear mixed effects model; Figure 6G). Heterozygous *ephA4b^ΔCLE/+^* mutant optic nerves were not significantly different from controls. Furthermore, eye size was not significantly different between conditions, eliminating delayed overall retinal growth as an explanation for the axon growth phenotype (wild-type N=30; *ephA4b^ΔCLE/+^* N=24; *ephA4b^ΔCLE/ΔCLE^* N=28; Figure 6H). This result demonstrates that complete loss of the *ephA4b-CLE* results in substantial reduction in RGC axon projection, indicating that the *ephA4b-CLE* is required for normal optic nerve outgrowth.

To further ascertain this phenotype, we employed an independent knockdown approach. We used a ‘splice morpholino’, which bound to the splice acceptor site of the *ephA4b-CLE,* and a mismatch morpholino with a mutated base as a control, validated previously^19^. Consistent with the genetic CLE deletion, embryos injected with the *ephA4b-CLE* splice morpholino displayed a normal retinal organisation (Supplementary Figure 7A-C) and an optic nerve volume loss compared with the uninjected and mismatch morpholino-injected controls (1.000 normalised volume in uninjected; N=20, 0.935 in mismatch-injected control; N=16, and 0.483 in morpholino-injected; N=17, P<0.005, linear mixed effects model; Supplementary Figure 7D-G). We also observed the same phenotype in F0 crispants, using the same CRISPR/Cas9 approach as above (1.000 normalised volume in uninjected; N=23, 1.191 in gRNA injected controls; N=29, 0.987 when a single RNP was injected; N=29, and 0.590 when both RNPs were injected, P<0.0001; N=32, linear mixed effects model Supplementary Figure 7H-L).

One way in which optic nerve volumes might be decreased is via reduced RGC axon growth rate. To test whether *ephA4b-CLE* LoF impacted RGC axon growth, the first RGC axon to leave the retina was visualised via timelapse confocal imaging (Figure 6I, J, Movie S1). We found that the mean RGC pioneer axon growth was significantly reduced in *ephA4b^ΔCLE/ΔCLE^*mutants (6.213µm/10minutes in wild-type; N=5, 4.180µm/10minutes in *ephA4b^ΔCLE/ΔCLE^*mutants; N=6, P<0.001 Welch’s test, Figure 6I – K, Movie S1). We comparably observed a similar decreased RGC pioneer axon growth in splice morpholino *ephA4b-CLE* LoF experiments (6.758µm/10minutes in uninjected; N=9, 7.229µm/10minutes in mismatch morpholino-injected; N=10, 3.248µm/10minutes in morpholino-injected; N=8, P<0.01 Kruskal-Wallis, Supplementary Figure 8A-D, Movie S2).

To test whether the optic nerve defect in *ephA4b^ΔCLE/ΔCLE^*mutants was a temporary issue, corrected later in development, we measured the volume of the nerve terminals in the optic tectum at 5dpf. This dorsal midbrain region receives the optic nerve endings and is required for visual processing. The size of its white matter is directly dependent on the neuronal projections connecting to tectal neurons^38,39^. We found that the volume of nerve endings in the 5dpf optic tectum was decreased in *ephA4b^ΔCLE/ΔCLE^* mutants compared with wild-types (1.043 normalised volume in wild-type vs 0.877 normalised volume in *ephA4b^ΔCLE/ΔCLE^*, Welch’s t test P<0.05, Figure 6L - N). This effect was also seen in morpholino experiments (1.000 normalised volume in uninjected wild-type, 1.144 normalised volume in mismatch morpholino-injected, 0.869 in morpholino-injected, Kruskal-Wallis, P<0.01, Supplementary Figure 8E-H).

Lastly, we sought to confirm that the *ephA4b-CLE* is indeed responsible for the RGC axonal growth defect, re-introducing expression of the *ephA4b-CLE* in the mutants. We did so by injecting a plasmid expressing *ephA4b-CLE-EGFP* under the control of the *rx3* promoter, which drives expression in the retinal progenitors and RGCs until ∼48hpf^40,41^, Figure 7A. This injection resulted in mosaic expression in the developing retinas during the formation of the optic nerves. We injected increasing doses of the plasmid into *ephA4b^ΔCLE/ΔCLE^* embryos and saw an increase in the volumes of the optic nerves with increasing doses (normalised to wild-type optic nerve volume 1.000 N=27: 0pg – 0.475, P<0.0001, N=26; 25pg – 0.698, P<0.05, N=25; 50pg – 0.789, P<0.0001, N=26; 75pg – 0.850, P<0.0001, N=29; 100pg – 1.004, P<0.0001, N=30; linear mixed effects model; Figure 7B-H). This rescue was not observed in uninjected embryos, showing that the optic nerve defect observed in the *ephA4b^ΔCLE/ΔCLE^* mutant is due to lack of CLE transcripts. These results confirmed that the optic nerve phenotype observed is caused by loss of *ephA4b-CLE* specifically.

**Figure 7:**
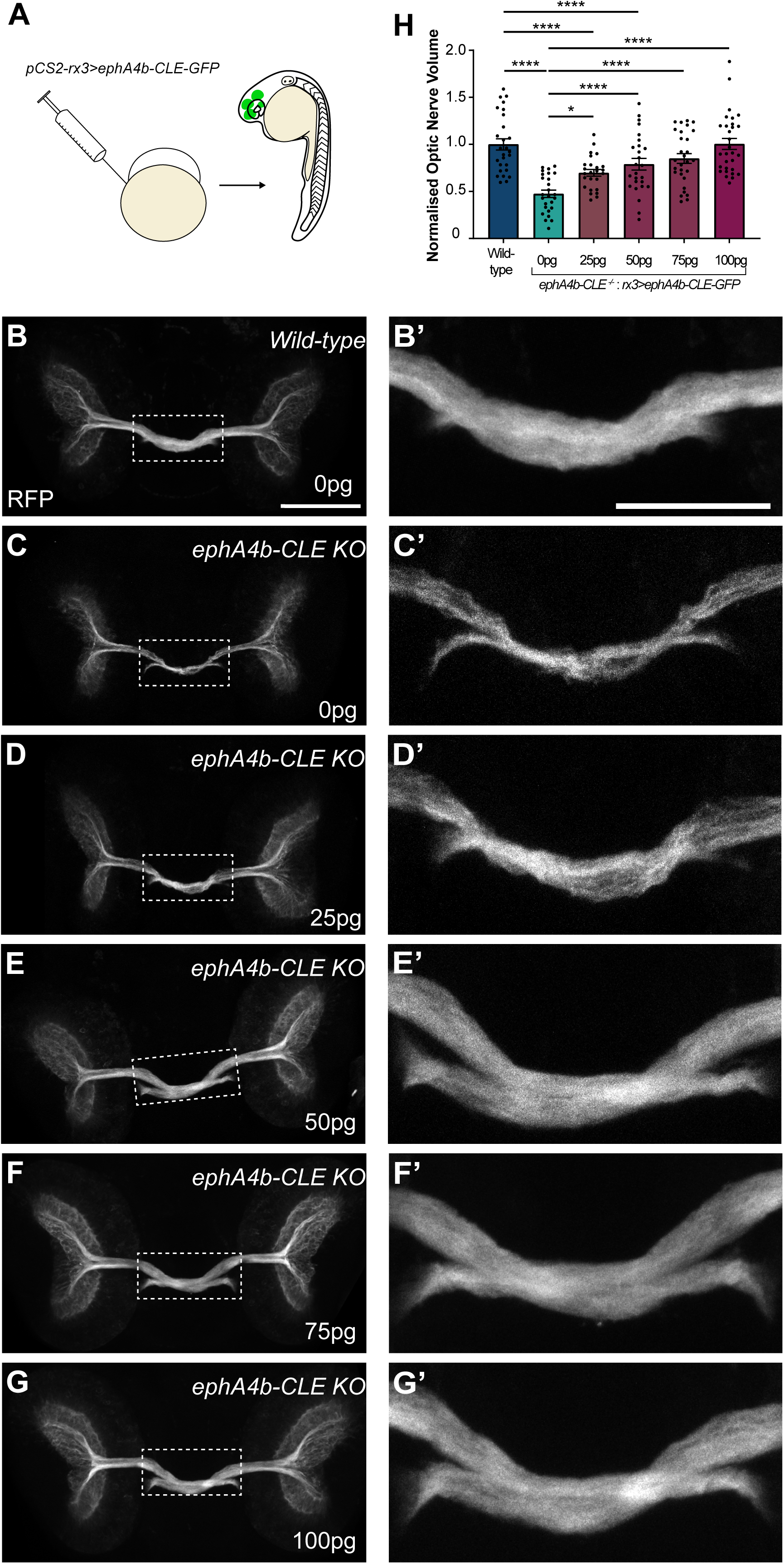
Re-expression of ephA4b-CLE rescues optic-nerve growth in ephA4b*^ΔCLE/ΔCLE^* mutants. **(A)** Schematic of the rescue strategy. *rx3:ephA4b-CLE-EGFP* plasmid was injected into one-cell-stage *ephA4b^ΔCLE/ΔCLE^* embryos to drive mosaic CLE expression in retinal progenitors and developing RGCs. **(B–G)** Maximum-intensity projections of 48-hpf *tg(ath5:RFP)* optic nerves from wild-type embryos (B) and *ephA4b^ΔCLE/ΔCLE^* embryos receiving 0 pg (C), 25 pg (D), 50 pg (E), 75 pg (F), or 100 pg (G) *rx3:ephA4b-CLE-EGFP*. Boxed regions are shown at higher magnification in B′–G′. **(H)** Quantification of normalised optic-nerve volume across the rescue series. Wild-type, N=27; mutant 0 pg, N=26; 25 pg, N=25; 50 pg, N=26; 75 pg, N=29; 100 pg, N=30. Increasing amounts of *rx3:ephA4b-CLE-EGFP* progressively restored optic-nerve volume, reaching approximately wild-type levels at 100 pg. Statistical significance was assessed using a linear mixed-effects model; *P<0.05, ****P<0.0001. Data are shown as mean ± SD. Scale bars = 50µm.

Loss of the CLE-containing interval abolishes *ephA4b-CLE* without measurably altering the assayed full-length transcript, causes a reproducible RGC axon-growth defect across stable genetic deletion and independent perturbation approaches, and the phenotype is rescued by re-expression of the CLE isoform. These experiments establish a requirement for *ephA4b-CLE* in normal RGC axon development. They do not establish whether the CLE acts by modulating full-length EphA4 abundance or signalling, and the downstream molecular mechanism remains to be determined.

Although CLE and competing splice usage show a modest inverse association across the broader CLE population, this relationship is not expected to produce stoichiometric changes in steady-state full-length RNA at every locus. At *ephA4b*, removal of the CLE abolished the CLE-containing isoform without significantly altering the abundance of the assayed full-length transcript, indicating that the developmental phenotype cannot be attributed simply to loss of total *ephA4b* expression.

### GAN-CLE regulates motor-axon extension and terminal branching.

Having established that loss of *ephA4b-CLE* disrupts optic nerve development, we next asked whether CLEs, not identified in disease-related datasets and associated with other developmentally relevant genes, also have functional roles in neuronal development. We chose as a case-study, a CLE located in intron 8 of *gigaxonin* (GAN). GAN has a defined role in zebrafish motor neuron development, where its protein promotes Sonic Hedgehog signalling to control motor neuron specification, axon pathfinding, and arborisation^42^. We detected GAN-CLE in our TRAP dataset and by PepQuery, indicating active ribosome association during early development. Using *tg(mnx1:RFP)* zebrafish to visualise motor neurons with RFP, we assessed whether disruption of *GAN-CLE* affected motor neuron development.

To disrupt GAN-CLE, we again used an F0 crispant approach, with two RNP complexes cutting either side of the CLE sequence (Figure 8A). Deletions were validated by PCR using genomic DNA at 28hpf and primers flanking the cut sites (425bp wild-type band, 185bp mutant band – red asterisk, Figure 8B). Mutant alleles were confirmed by sequencing. Increasing concentrations of RNPs were injected, from a 1:64 to 1:2 dilution of the initial 30.5µM RNP mix, to identify a concentration that produced efficient deletion, while minimising editing products. The 1:8 dilution produced a clear wild-type and deletion-specific band and was therefore used for subsequent experiments (Figure 8B).

**Figure 8:**
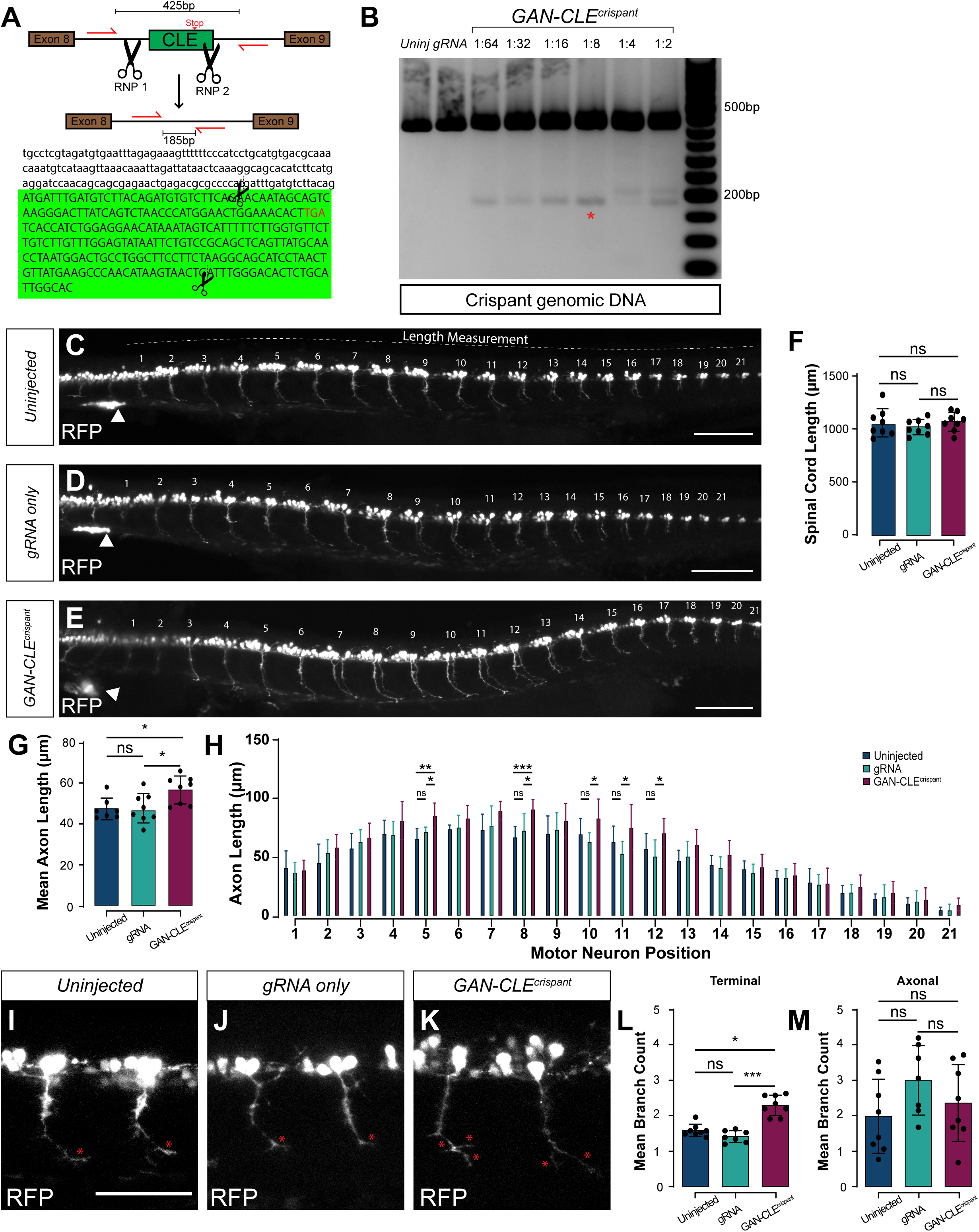
Disruption of GAN-CLE increases motor-axon extension and terminal branching. **(A)** Schematic of the F0 CRISPR/Cas9 strategy targeting GAN-CLE. Two RNP complexes flanking the CLE remove an approximately 240-bp region of *GAN* intron 8; wild-type and deletion alleles generate approximately 425-bp and 185-bp PCR products, respectively. **(B)** PCR validation of GAN-CLE disruption in 28-hpf embryos across uninjected, gRNA-only, and increasing RNP concentrations from 1:64 to 1:2 of the initial 30.5-µM RNP mixture. The 1:8 condition was used for phenotypic analysis; the deletion-specific band is indicated by the red asterisk. **(C–E)** Maximum-intensity projections of *tg(mnx1:RFP)* motor neurons in uninjected (C), gRNA-only (D), and *GAN-CLE^crispant^* (E) embryos at 28 hpf. Motor-neuron positions 1–21 are indicated. White arrowheads mark *tg(mnx1:RFP)*-positive anatomical landmarks used for spinal-cord-length measurements. **(F)** Spinal-cord length was not significantly different between conditions (N=8 embryos per group; one-way ANOVA). **(G)** Mean motor-axon length per embryo was increased in *GAN-CLE^crispant^* embryos compared with controls (N=8; one-way ANOVA with Tukey’s multiple comparisons, *P<0.05). **(H)** Motor-axon length across motor-neuron positions 1–21. Statistical significance was assessed using a two-way linear mixed-effects model with embryo as the repeated-measures unit followed by Šídák-adjusted comparisons within motor-neuron position; *P<0.05, **P<0.01, ***P<0.001, ns, not significant. **(I–K)** Representative motor neurons from uninjected (I), gRNA-only (J), and *GAN-CLE^crispant^* (K) embryos. Red asterisks indicate terminal axonal branches. **(L–M)** Mean numbers of terminal branches (L) and non-terminal axonal branches (M) per embryo. GAN-CLE disruption increased terminal branching, whereas non-terminal branch number was unchanged; Kruskal–Wallis test with Dunn’s multiple comparisons, *P<0.05, ***P<0.001, ns, not significant. Data are shown as mean ± SD. Scale bars = 50µm.

We examined whether loss of *GAN-CLE* altered motor neuron development in *tg(mnx1:RFP)* embryos at 28hpf. Motor neurons were visualised by confocal microscopy in uninjected, gRNA-only-injected and *GAN-CLE^crispant^*embryos (Figure 8C–E). To determine if there were differences in embryo size or developmental progression, we measured spinal cord length using mnx1:RFP-positive endodermal cells (Figure 8C-E, white arrowhead) as anatomical landmarks (representative length measurement dotted line in Figure 8C). Spinal cord length was not significantly different across conditions (Figure 8F, N=8 embryos, one-way ANOVA).

We next quantified axon length across motor neuron positions 1-21 (Figure 8C-E, G-H). *GAN-CLE^crispant^* embryos showed an overall mean increase in motor axon length compared to uninjected and gRNA-only-injected controls (uninjected = 47.4µm; gRNA-only-injected = 47.7µm; *GAN^crispant^*= 56.6µm; N=8 embryos, P<0.05, one-way ANOVA with Tukey’s multiple comparisons, Figure 8G). Axon length across individual motor neuron positions displayed a general trend of increased lengths in *GAN-CLE^crispant^* embryos, however only reached significance in a specific motor neuron positions (5, 8, 10, 11, 12, N=8, *** = P<0.001 compared with uninjected for positions 5 and 8; * = P<0.05 compared with gRNA for all each condition, two-way linear mixed effects model with post hoc comparisons performed using Šídák-adjusted multiple comparisons, Figure 8H). Overall, this shows that motor axon extension is enhanced by loss of GAN-CLE, indicating a role for the CLE mRNA expression in controlling motor axon growth and/or maturation.

To assess axonal maturation, we measured the number of branches along each motor axon and the number of terminal branches. In both cases, only branches that were longer than 5µm were counted to distinguish from transient filopodia. We found that *GAN-CLE^crispant^*embryos had more terminal branches compared to uninjected and gRNA-only-injected controls (uninjected = 1.58 branches, gRNA-only-injected = 1.41 branches, *GAN-CLE^crispant^* embryos = 2.28 branches, * = P<0.05, *** = P<0.001, N=8, Kruskal-Wallis test with Dunn’s multiple comparisons, terminal branches labelled with red asterisk, Figure 8I-L). Contrastingly, we observed that the mean number of non-terminal axonal branches per embryo was not significantly different across conditions (Figure 8M), pointing to an acceleration of axonal maturation.

Together these results show that *GAN-CLE* is required for normal motor neuron morphology and axonal maturation. Unlike loss of *ephA4b-CLE*, which reduced RGC axonal growth, *GAN-CLE* disruption increased motor axon extension and terminal branching. Since GAN has been shown to regulate Sonic Hedgehog signalling in motor neurons, which drives axonal growth and branching^42^, GAN-CLE may act as a negative regulator of the full-length gene, the disruption of which would promote Sonic Hedgehog signalling and motor axon growth.

Because F0 CRISPR disruption can produce mosaic editing and potentially locus-independent effects, we sought to validate the GAN-CLE phenotype using an independent perturbation strategy. We designed a splice-blocking morpholino targeting GAN-CLE and first assessed its molecular specificity by RT-qPCR. GAN-CLE expression was strongly reduced across a morpholino dose series, to 0.196, 0.123 and 0.050 relative expression at 1:20, 1:10 and 1:5 dilutions, respectively, compared with 1.000 in uninjected embryos (ordinary one-way ANOVA, P=3.85×10⁻⁷; all morpholino doses versus uninjected, Tukey-adjusted P<7×10⁻⁶; Supplementary Figure 9A). In contrast, full-length *GAN* transcript abundance was not significantly altered (ordinary one-way ANOVA, P=0.063; Supplementary Figure 9B), demonstrating selective disruption of the CLE-containing isoform.

We then examined motor-neuron morphology following GAN-CLE splice blocking in *tg(mnx1:RFP)* embryos. As in the F0 crispants, overall spinal-cord length was unchanged (uninjected, 1314.9 µm; mismatch morpholino, 1311.2 µm; GAN-CLE morpholino, 1296.2 µm; N=7, 5 and 9 embryos respectively; one-way ANOVA, P=0.786; Supplementary Figure 9C–F), whereas mean motor-axon length was significantly increased (uninjected, 72.4 µm; mismatch morpholino, 70.8 µm; GAN-CLE morpholino, 84.0 µm; one-way ANOVA, P=0.0029; Tukey-adjusted GAN-CLE morpholino versus uninjected P=0.0097 and versus mismatch P=0.0079; Supplementary Figure 9G). Thus, an independent splice-blocking perturbation selectively reducing GAN-CLE recapitulated the increased motor-axon extension observed following CRISPR disruption, substantially reducing the likelihood that the F0 phenotype results from CRISPR-specific or off-target effects.

## Discussion

CLEs were initially characterised in settings of RBP dysfunction and neurodegenerative disease, where they were interpreted primarily as aberrant splice products^19,43–45^. Our data substantially broaden this view. CLE-containing RNAs are detectable across physiological zebrafish, mouse and human transcriptomes, a subset is developmentally regulated, and CLE usage is unusually sensitive to perturbation of neural RBPs. Some CLE transcripts associate with ribosomes, CLE-specific peptides can be detected for defined candidates, and endogenous tagging demonstrates that *ephA4b-CLE* produces a stable truncated protein during normal development. Together with the developmental phenotypes caused by loss of *ephA4b-CLE* and *GAN-CLE*, these findings establish that CLEs are not restricted to pathological splicing failure, but form part of the physiological transcript repertoire.

This does not imply that every CLE is functional, in fact most physiological CLEs are low-usage events. Furthermore, only a subset show significant developmental regulation and ribosome association alone does not demonstrate stable protein production. CLEs may therefore span a continuum from low-frequency terminal-splicing events with little biological consequence to regulated isoforms with specific RNA- or protein-level functions. Importantly, low abundance cannot itself be taken as evidence of noise: *ephA4b-CLE* is a minor physiological isoform yet generates a detectable endogenous protein and is required for normal RGC development. This distribution is reminiscent of the evolutionary behaviour of newly created exons, which are commonly incorporated at low frequency and remain alternatively spliced, allowing a new transcript isoform to coexist with the ancestral product while exposing it to subsequent regulatory and functional selection^46–49^.

The evolutionary properties of CLEs suggest that their recurrence is driven more strongly by genomic context than by conservation of the cryptic exon sequence itself. CLEs show little primary-sequence conservation, align poorly across species and retain the transposable-element characteristics of their surrounding introns. Nevertheless, CLE-containing host genes recur across vertebrates more frequently than expected, and a subset of CLEs occupy orthologous introns. Together with their enrichment in earlier, intermediate-length introns, these observations suggest that particular genes and intronic environments are permissive for CLE formation, potentially through combinations of splice sites, polyadenylation signals and RBP-binding elements.

These properties place CLEs within the broader process of exonisation, in which previously intronic sequence becomes incorporated into mature transcripts. Transposable elements are an important substrate for exon birth because they can harbour splice-recognition and 3′-end-processing signals, and TE-derived exons are typically used as alternative rather than constitutive products^50–53^. CLEs may therefore represent a terminal-exon form of exonisation, in which recognition of an intronic splice acceptor is coupled to cleavage and polyadenylation to create a new transcript end. Consistent with this model, CLEs retain an intron-like TE composition rather than resembling conventional exons. Functional precedents exist and include a LINE-1-derived terminal exon in *ATRN* that generates a soluble Attractin isoform and an Alu-derived terminal exon in *IFNAR2* that produces a physiological truncated decoy receptor^54,55^.

Intriguingly, CLEs recurring in orthologous host genes contain less recognisable TE sequence and are more frequently TE-free than species-specific CLEs. This may indicate that TEs disproportionately seed evolutionarily young, lineage-restricted CLEs, whereas recurrent CLE formation is more often supported by TE-independent intronic architecture. Alternatively, TE-derived CLEs may progressively diverge during exonisation such that their ancestral repeat identity becomes difficult to recognise. Our data cannot distinguish between these possibilities but suggest that TE sequence may contribute more strongly to the generation of CLE diversity than to the long-term maintenance of recurrent CLE usage.

Across 20 RBP perturbation comparisons, CLE responses were bidirectional and RBP-specific, but CLE usage was consistently more labile than that of matched conventional ALEs and generally changed more strongly than the dominant terminal exon from the same gene. CLE formation therefore appears to sit at a sensitive boundary between cryptic-site repression and productive terminal-exon definition. hnRNP C suppresses recognition of thousands of cryptic Alu splice sites by competing with U2AF at polypyrimidine tracts, and loss of this repression exposes previously cryptic exons^56^. Loss of SFPQ, TDP-43 or related factors may amplify splice choices that already exist within the physiological repertoire rather than necessarily creating entirely novel transcripts. Disease may therefore shift the quantitative position of an existing cryptic-to-productive splice equilibrium, turning a normally rare or spatially restricted product into a dominant transcript. This interpretation provides a potential bridge between the physiological CLE collection identified here and the dramatic CLE activation observed following disruption of neuronal splicing factors.

The *EphA4* locus provides a striking example of how evolutionary flexibility can nevertheless converge on a similar functional output. Human *EPHA4* and both zebrafish *ephA4* paralogues contain CLEs within the orthologous intron downstream of the same conserved coding exon, despite no detectable homology between the CLE sequences. These independently arising isoforms truncate EphA4 at a similar position, retaining the ephrin-binding domain while removing the transmembrane and intracellular signalling regions, and AlphaFold3 modelling predicts preservation of the ephrin-binding interface. This architecture is notable given that naturally occurring truncating *EPHA4* variants have been reported in cancer, ALS and neurodevelopmental disease, including p.Arg230Ter within the cysteine-rich ectodomain, emphasising the functional importance of truncation position^57–60^. There is also precedent for regulating EphA4 by separating its extracellular and signalling domains. EphA4 undergoes developmentally regulated ectodomain shedding, while soluble EphA4 ectodomains can antagonise Eph/ephrin signalling and promote neurite or axonal growth^61,62^. In retinal ganglion cells, reduced EphA4 forward signalling similarly increases axon extension^63^. Together with the retained ephrin-binding interface predicted here, these observations suggest that EphA4b-CLE could act as a soluble or signalling-deficient ligand-binding isoform. This would parallel *IFNAR2*, where exonisation of intronic sequence generates a physiological truncated decoy receptor^54^. However, secretion, ligand binding and competitive antagonism by EphA4b-CLE remain to be demonstrated experimentally.

The functional data establish biological importance of the zebrafish *ephA4b* CLE independently of this mechanistic model. By contrast, disruption of *GAN-CLE* increased motor-axon extension and terminal branching and was reproduced by an independent splice-blocking morpholino. These opposite phenotypes argue that CLEs provide a route to transcript innovation rather than a common downstream mechanism: their effects depend on the host gene, protein domains retained or removed, and properties of the resulting RNA or protein.

These findings also reshape the interpretation of disease-associated CLEs since those mis-regulated by RBP dysfunction may represent quantitative or contextual dysregulation of normally accessible splice outcomes rather than exclusively pathological transcripts, arguing for event-specific rather than indiscriminate suppression of cryptic splicing. Their strong responsiveness to neural RBP state may also make CLE junctions useful molecular readouts of splicing-network dysfunction, although their clinical utility remains to be established. Overall, our results identify CLEs as a physiological source of vertebrate transcript diversity.

## Supporting information

Supplementary Datasheet 1

Supplementary Datasheet 2

Supplementary Datasheet 3

Supplementary Datasheet 4

Supplementary Datasheet 5

Supplementary Datasheet 6

Supplementary Datasheet 7

Supplementary Datasheet 8

Supplementary Datasheet 9

Supplementary Datasheet 10

Supplementary Datasheet 11

## Acknowledgments

We thank Brant Weinstein for the generous gift of TRAP-expressing fish which enabled us to make the ubiquitous TRAP-expressing plasmid. We also acknowledge Chintan Trivedi for help and advice with hybridisation chain reaction. This work is supported by funds from the Wellcome Trust (WT 220861/Z/20/Z to CH).

Conceptualisation: **MPB**, CH

Investigation: **MPB**, KLG, FH, LOA, CL

Supervision: **MPB**, CH, FH

Funding: CH, **MPB**

Writing – original draft: **MPB**

Writing – review and editing: **MPB**, FH, KLG, CH

---

**Supplementary Figure 1:**
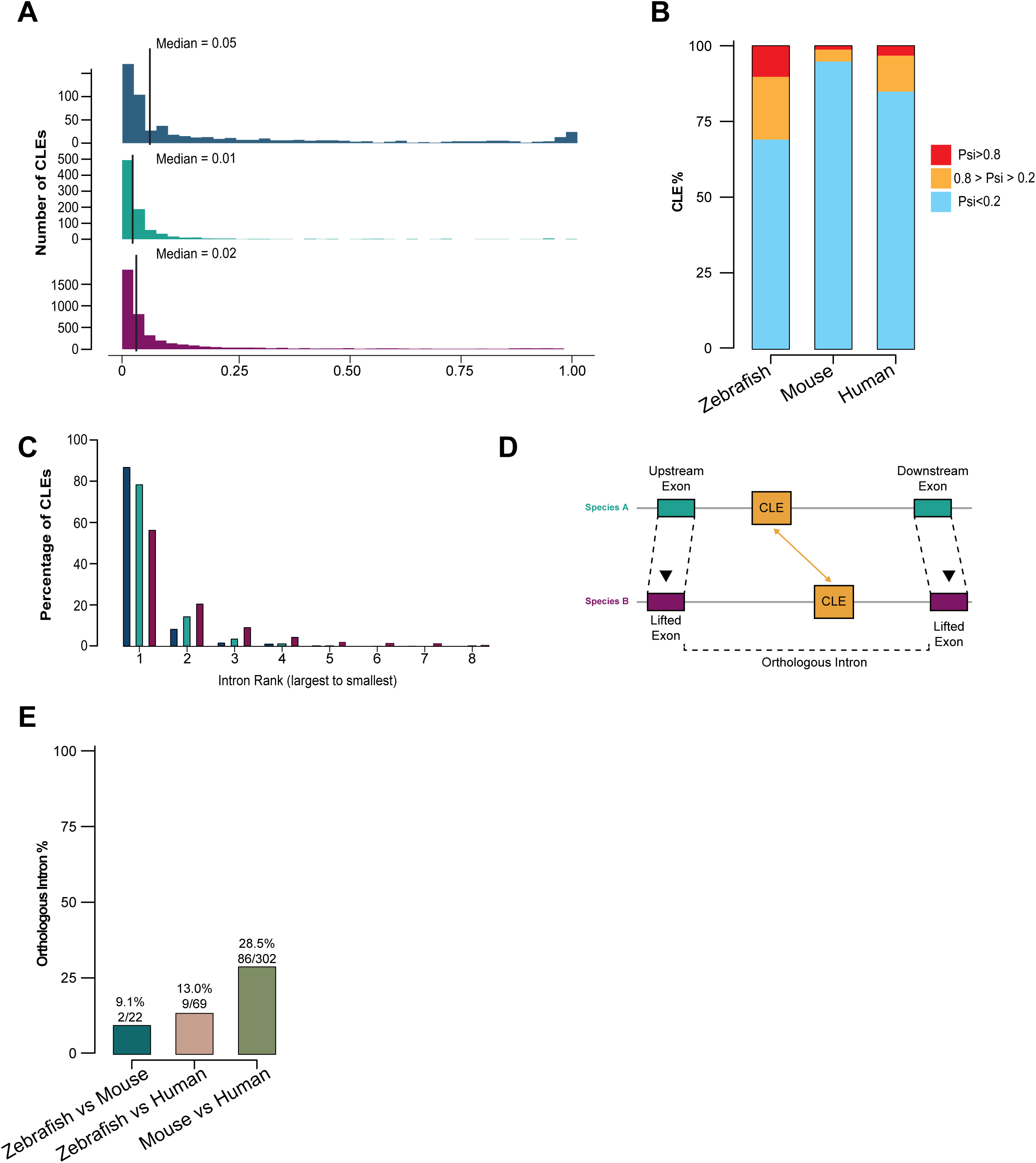
Physiological CLEs are predominantly low-usage isoforms and show limited positional conservation across species. **(A)** Distribution of CLE PSI in zebrafish, mouse and human, calculated from splice-junction reads arising from the shared upstream donor. Median PSI was 0.05 in zebrafish, 0.01 in mouse and 0.02 in human. **(B)** Percentage of CLEs with low (PSI<0.2), intermediate (0.2≤PSI≤0.8), or high (PSI>0.8) inclusion in each species. CLEs predominantly represented low-usage transcript isoforms. **(C)** Distribution of CLE-containing introns according to intron size rank within each host gene, where rank 1 represents the longest intron. CLEs occurred within the longest intron for 87.4% of zebrafish, 79.0% of mouse and 57.0% of human events. **(D)** Schematic of the orthologous-intron analysis. Exons immediately upstream and downstream of a CLE-containing intron were projected into the partner-species genome by liftOver; a CLE in the interval flanked by the corresponding projected exons was classified as occupying an orthologous intron. **(E)** Percentage of testable shared-gene CLEs occupying orthologous introns. Orthologous-intron positioning was observed for 2/22 zebrafish–mouse comparisons (9.1%), 9/69 zebrafish–human comparisons (13.0%) and 86/302 mouse–human comparisons (28.5%).

**Supplementary Figure 2:**
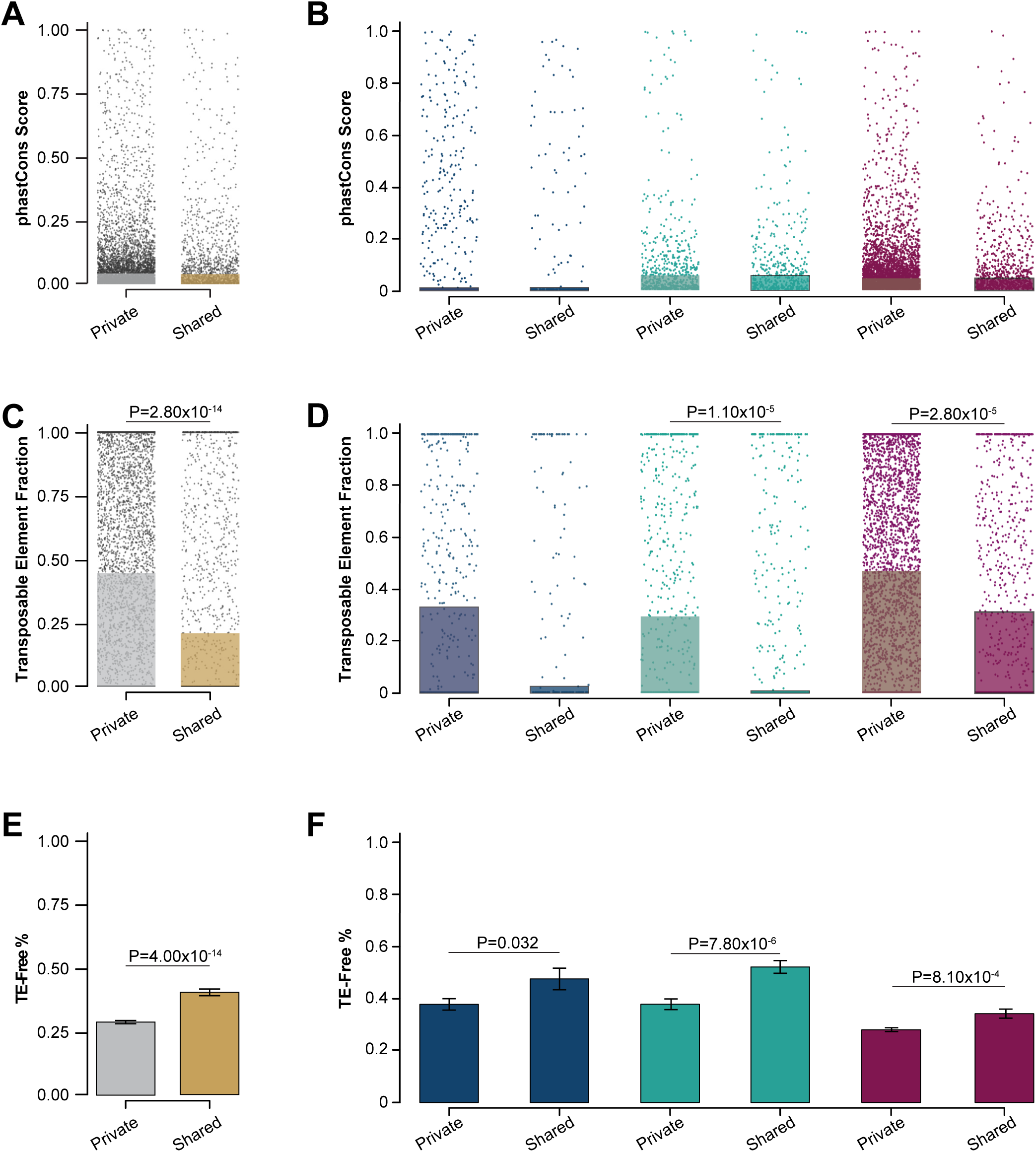
CLE recurrence across orthologous genes is not associated with increased sequence conservation but is associated with reduced transposable-element content. **(A)** phastCons conservation scores of species-specific (“private”) CLEs compared with CLEs whose host gene contained a CLE in at least one additional species (“shared”). Shared CLEs were not more conserved than private CLEs (median phastCons 0.039 versus 0.038; two-sided Wilcoxon rank-sum test, not significant). **(B)** Species-resolved phastCons scores for private and shared CLEs in zebrafish, mouse and human; no significant increase in conservation was detected for shared CLEs in any species. **(C)** Transposable-element (TE) fraction of pooled private and shared CLEs. Shared CLEs contained less TE-derived sequence than private CLEs (median TE fraction 0.21 versus 0.44; two-sided Wilcoxon rank-sum test, P=2.8×10⁻¹⁴). **(D)** Species-resolved TE fractions for private and shared CLEs. Shared CLEs showed lower TE content in mouse (P=1.1×10⁻⁵) and human (P=2.8×10⁻⁵), with the same directional trend in zebrafish; two-sided Wilcoxon rank-sum tests. **(E)** Proportion of pooled CLEs containing no annotated TE sequence. Shared CLEs were more frequently TE-free than private CLEs (41.6% versus 30.2%; Fisher’s exact test, P=4.0×10⁻¹⁴). **(F)** Species-resolved proportions of TE-free CLEs. TE-free events were more frequent among shared CLEs in zebrafish (47.9% versus 37.7%, P=0.032), mouse (52.5% versus 37.9%, P=7.8×10⁻⁶) and human (34.3% versus 28.0%, P=8.1×10⁻⁴); Fisher’s exact tests.

**Supplementary Figure 3:**
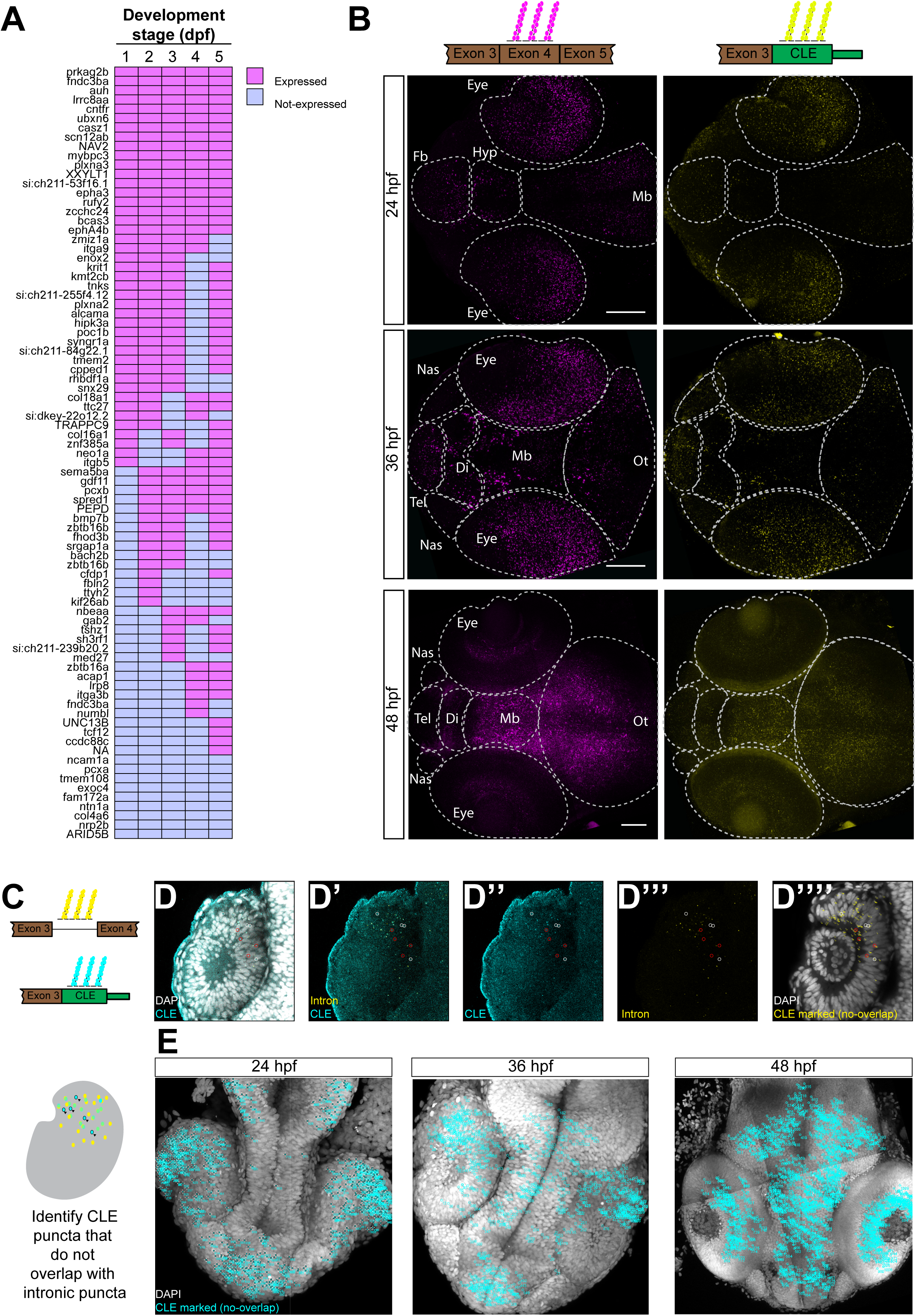
SFPQ-sensitive CLEs are expressed during normal zebrafish development and ephA4b-CLE-region signal is detected in developing neural tissues. **(A)** Developmental RT-PCR detection summary for SFPQ-sensitive CLEs whose primer pairs were independently validated using *sfpq*^-/-^ positive-control cDNA. CLEs were analysed in wild-type zebrafish at 1, 2, 3, 4 and 5 dpf. Magenta indicates detection of the expected CLE-containing product and blue indicates no detectable product. Of 83 technically validated assays, 74 CLEs (89.2%) were detected during at least one wild-type developmental stage and nine were not detected under the assay conditions. **(B)** Hybridisation Chain Reaction (HCR) analysis of the *ephA4b* locus at 24, 36 and 48 hpf using probes targeting exon 4 of the full-length transcript (magenta; left) and the CLE-containing region (yellow; right). Dorsal views are shown with anatomical regions outlined. CLE-region signal was detected within the broader *ephA4b* expression domain, including developing neural structures. Fb, forebrain; Hyp, hypothalamus; Tel, telencephalon; Di, diencephalon; Nas, nasal pits; Mb, midbrain; Ot, optic tectum. **(C)** Schematic of the HCR strategy used to distinguish puncta arising from the CLE region from signal overlapping the surrounding host intron. Probes targeting the CLE sequence are shown in cyan and probes targeting the intron containing the CLE are shown in yellow. CLE puncta that did not overlap with intronic puncta were scored as signal consistent with the CLE-containing transcript. **(D)** Representative HCR analysis of *ephA4b-CLE* in a single optical section through a 24 hpf zebrafish retina. CLE-targeting probes are shown in cyan, probes targeting the downstream host intron in yellow, and DAPI in grey. **(D)** Merged CLE, intronic and DAPI channels. White circles indicate CLE-positive puncta overlapping DAPI-positive nuclear regions, whereas red circles indicate CLE-positive puncta spatially separated from DAPI-positive nuclei and lacking overlapping intronic signal. **(D’)** CLE and intronic probe channels. **(D’’)** CLE channel alone. **(D’’’)** intronic-probe channel alone. **(D’’’’)** DAPI maximum projection showing the positions of CLE-specific puncta identified across the corresponding optical sections. **(E)** DAPI maximum projections of zebrafish retinas at 24, 36 and 48 hpf showing the positions of CLE-specific puncta identified across confocal z-stacks. Marked puncta represent CLE-probe signal lacking coincident downstream-intronic signal; red circles indicate CLE-positive puncta spatially separated from DAPI-positive nuclear regions and white circles indicate CLE-positive puncta overlapping DAPI-positive nuclei. Scale bars = 100µm.

**Supplementary Figure 4:**
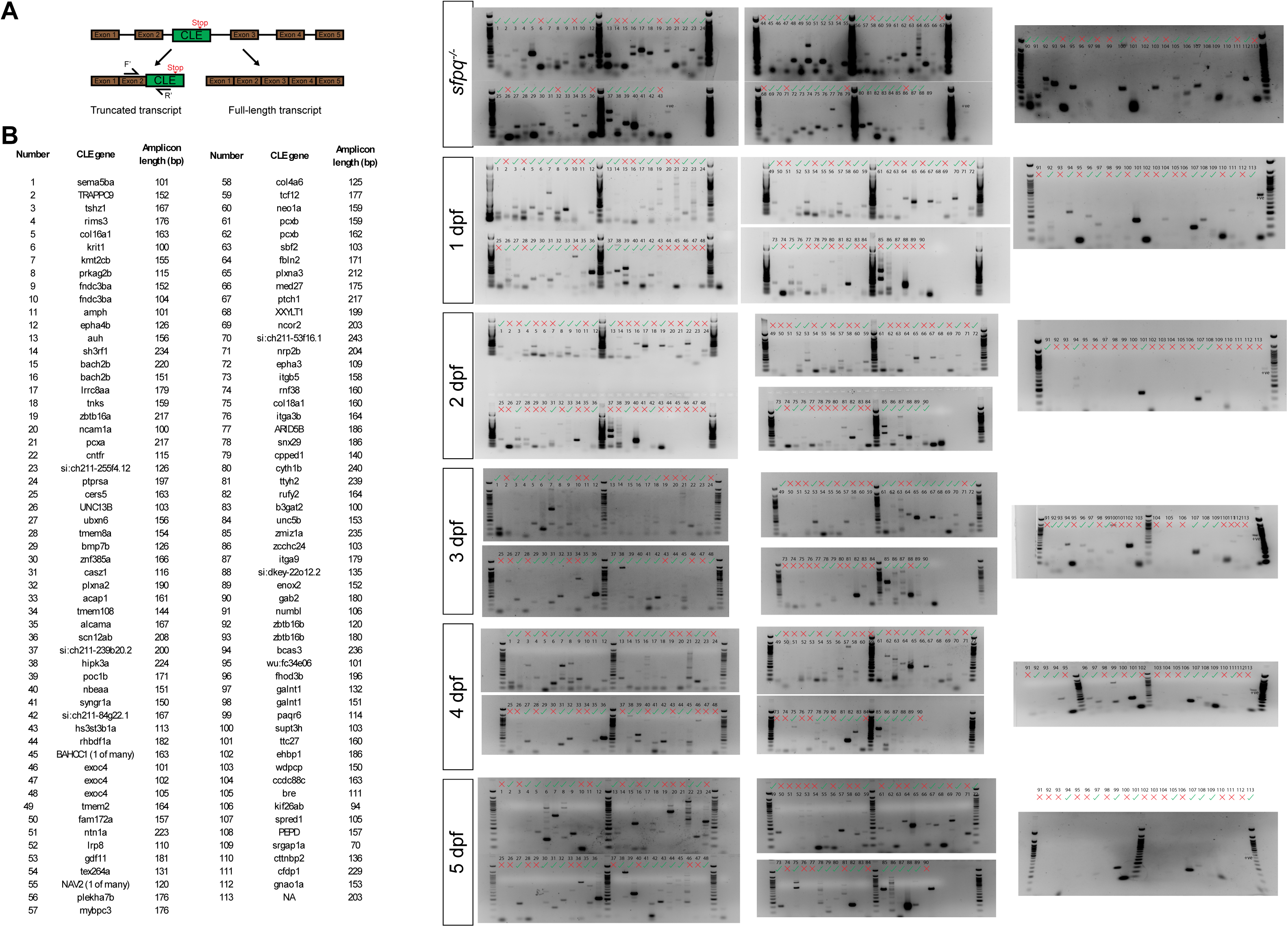
RT-PCR validation of SFPQ-sensitive CLE expression during normal zebrafish development. (A) Schematic of the CLE-specific RT-PCR strategy. Forward primers were positioned in the annotated exon immediately upstream of the CLE and reverse primers within the CLE, generating an amplicon across the upstream exon-to-CLE splice junction. (B) RT-PCR screen of 113 CLEs previously identified following loss of SFPQ. Primer performance was independently assessed using *sfpq*^-/-^ cDNA as a positive control, followed by analysis of wild-type zebrafish RNA at 1, 2, 3, 4 and 5 dpf. CLE numbers correspond to the accompanying gene and expected-amplicon-size table. Green ticks indicate detection of a product at the expected size and red crosses indicate no expected product. Eighty-three of 113 assays generated the expected product in *sfpq*^-/-^ cDNA and were therefore considered technically validated; 74/83 of these CLEs were detected during at least one wild-type developmental stage. Assays failing the *sfpq*^-/-^ positive control were considered technically uninformative and were not scored as absent from wild-type animals.

**Supplementary Figure 5:**
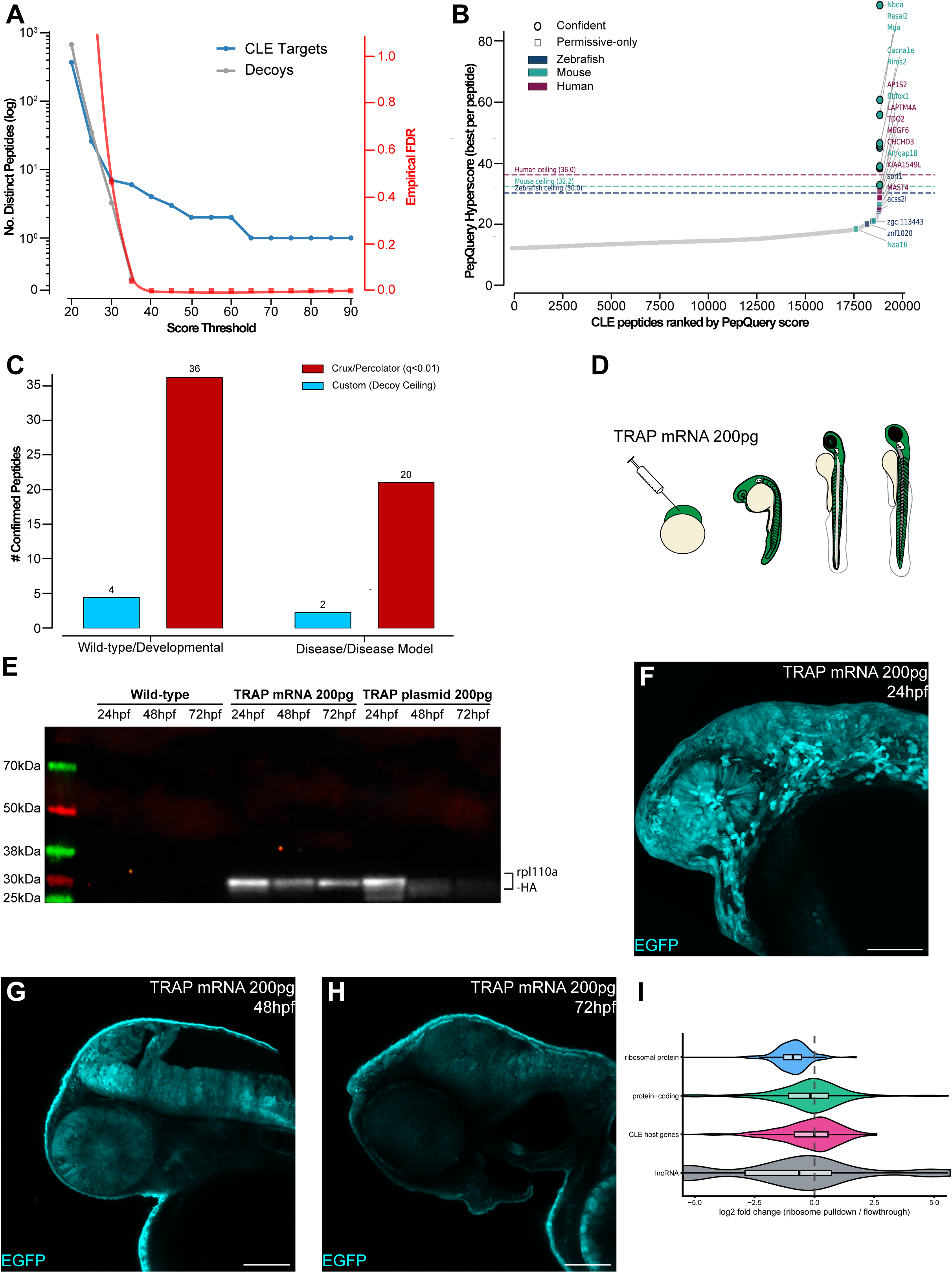
Proteomic search validation and characterisation of the TRAP system. **(A)** Relationship between PepQuery score threshold, numbers of distinct CLE target and composition-matched decoy peptides detected, and the corresponding empirical false-match rate. Increasing score thresholds progressively reduced the decoy background. **(B)** CLE-specific peptides ranked by maximum PepQuery hyperscore across the public proteomic datasets. Horizontal dashed lines indicate the maximum score attained by any of the eight composition-matched decoy peptide sets for zebrafish, mouse and human. Peptides are distinguished according to whether they passed the stringent species-wide decoy ceiling or were retained only by the more permissive search criteria. **(C)** Number of CLE-specific peptides detected in wild-type/developmental and disease/disease-model proteomic datasets using the composition-matched decoy-ceiling approach or Crux/Percolator at peptide q<0.01. **(D)** Schematic of the zebrafish TRAP experiment. One-cell-stage embryos were injected with 200 pg *rpl10a-3xHA-P2A-EGFP* mRNA and analysed across the first three days of development. **(E)** Anti-HA western blot validating expression of Rpl10a-3xHA following injection of 200 pg TRAP mRNA or plasmid. Wild-type embryos were used as negative controls. Expression following mRNA injection remained detectable at 24, 48 and 72 hpf. **(F–H)** Representative EGFP fluorescence in embryos injected with 200 pg TRAP mRNA at 24 hpf (F), 48 hpf (G) and 72 hpf (H), confirming persistence of construct expression during the TRAP collection period. **(I)** Gene-level partitioning of transcripts between the ribosome pull-down and flow-through fractions (DESeq2 log₂ fold change, pull-down/flow-through). CLE-host genes partition with translated protein-coding mRNA and are significantly distinct from non-coding lncRNAs, which are depleted from the ribosome fraction (Wilcoxon P = 2×10⁻⁴); ribosomal-protein mRNAs shown for reference. Scale bars = 100µm.

**Supplementary Figure 6:**
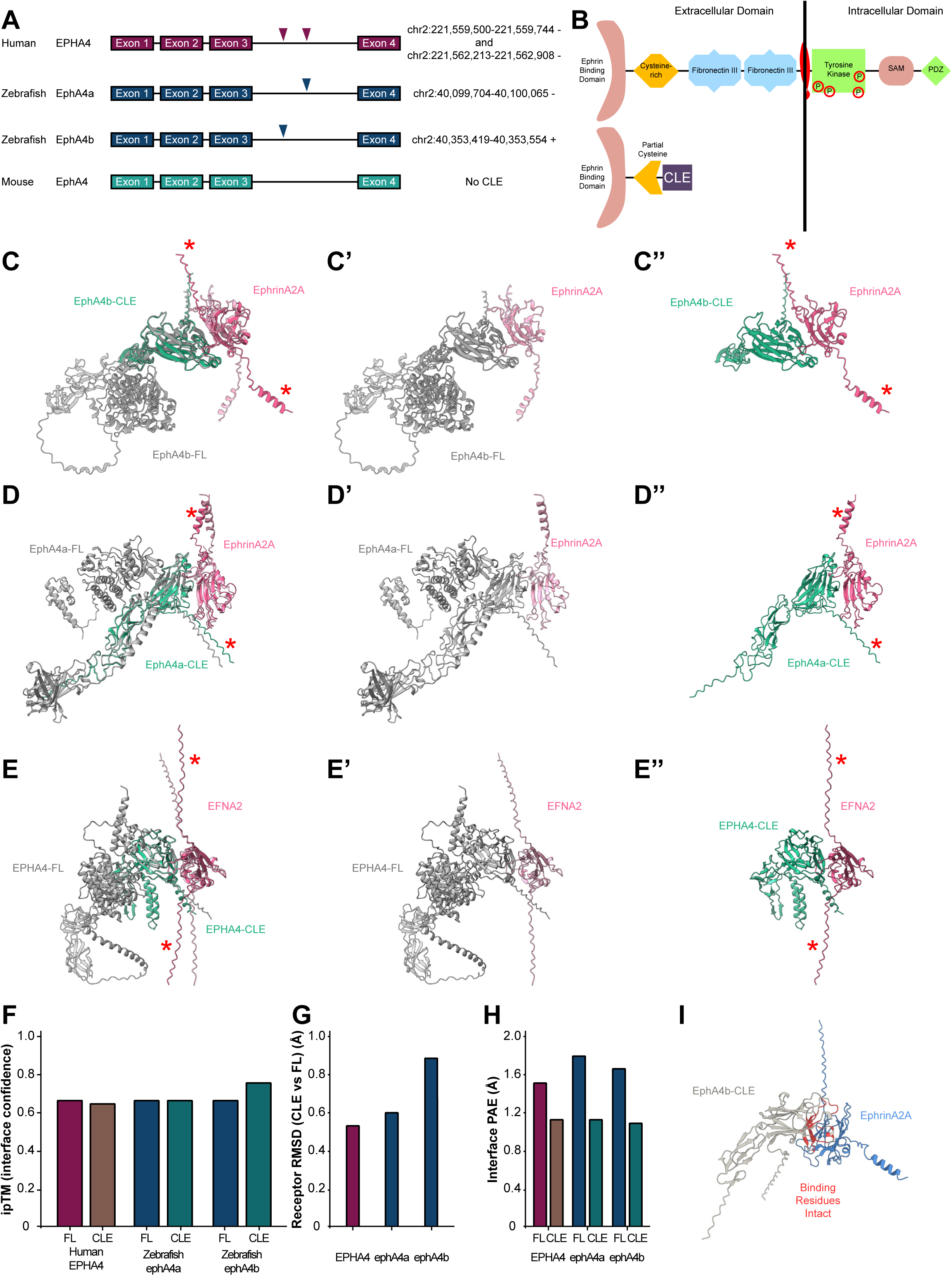
Convergent EphA4 cryptic last exons generate predicted soluble ectodomains with preserved ephrin-binding interfaces. **(A)** Gene architecture of *EphA4* orthologues aligned across human, mouse and zebrafish. Two CLEs were identified in human *EPHA4*, and one CLE each in zebrafish *ephA4a* and *ephA4b*. In each case, the CLE arises within the orthologous intron downstream of the same conserved coding exon. No equivalent CLE was detected in mouse *Epha4*. CLE genomic coordinates and transcriptional orientation are indicated. **(B)** Domain architecture of full-length EphA4 and the predicted CLE-derived protein. Full-length EphA4 contains an N-terminal ephrin-binding domain, cysteine-rich region, two fibronectin type III repeats, transmembrane segment, intracellular tyrosine kinase domain, SAM domain and C-terminal PDZ-binding motif. CLE inclusion terminates translation within the extracellular region, retaining the complete ephrin-binding domain and part of the cysteine-rich region while removing both fibronectin repeats, the transmembrane segment and all intracellular signalling domains. **(C–C′′)** AlphaFold3 models of zebrafish EphA4b in complex with ephrinA2a. **(C)** Superposition of full-length EphA4b (grey) and EphA4b-CLE (teal) receptor models with their respective predicted ephrinA2a complexes. **(C′)** Full-length EphA4b–ephrinA2a complex. **(C′′)** EphA4b-CLE–ephrinA2a complex. Asterisks indicate the ligand C-terminal membrane-anchoring region. **(D–D′′)** Corresponding AlphaFold3 models of zebrafish EphA4a and EphA4a-CLE in complex with ephrinA2a. **(E–E′′)** Corresponding AlphaFold3 models of human EPHA4 and a representative human EPHA4 CLE in complex with human EFNA2. **(F)** AlphaFold3 interface predicted TM-score (ipTM) for full-length and CLE receptor–ephrin complexes. CLE complexes showed interface-confidence values comparable to the corresponding full-length complexes: human EPHA4, 0.66 versus 0.68; zebrafish EphA4a, 0.69 versus 0.69; and zebrafish EphA4b, 0.75 versus 0.69. **(G)** Backbone RMSD between CLE and full-length receptor ectodomains following structural superposition. RMSD values were below 1 Å for human EPHA4 (0.54 Å), zebrafish EphA4a (0.60 Å) and zebrafish EphA4b (0.87 Å), indicating close structural agreement between the retained CLE-derived region and the corresponding region of the full-length receptor. **(H)** Mean interface predicted aligned error (PAE) for full-length and CLE receptor–ephrin complexes. CLE complexes showed lower predicted interface uncertainty than their corresponding full-length models in human EPHA4 (1.13 versus 1.50 Å), zebrafish EphA4a (1.19 versus 1.76 Å) and zebrafish EphA4b (1.09 versus 1.64 Å). **(I)** AlphaFold3 model of the zebrafish EphA4b-CLE–ephrinA2a complex showing receptor residues predicted to contact ephrinA2a within 5Å (red). All identified receptor-contact residues lie within the retained ligand-binding domain, indicating preservation of the predicted ephrin-binding interface following CLE-mediated truncation. Full AlphaFold3 confidence metrics, interface measurements, sequences and structural comparisons are provided in Supplementary Datasheet 11.

**Supplementary Figure 7:**
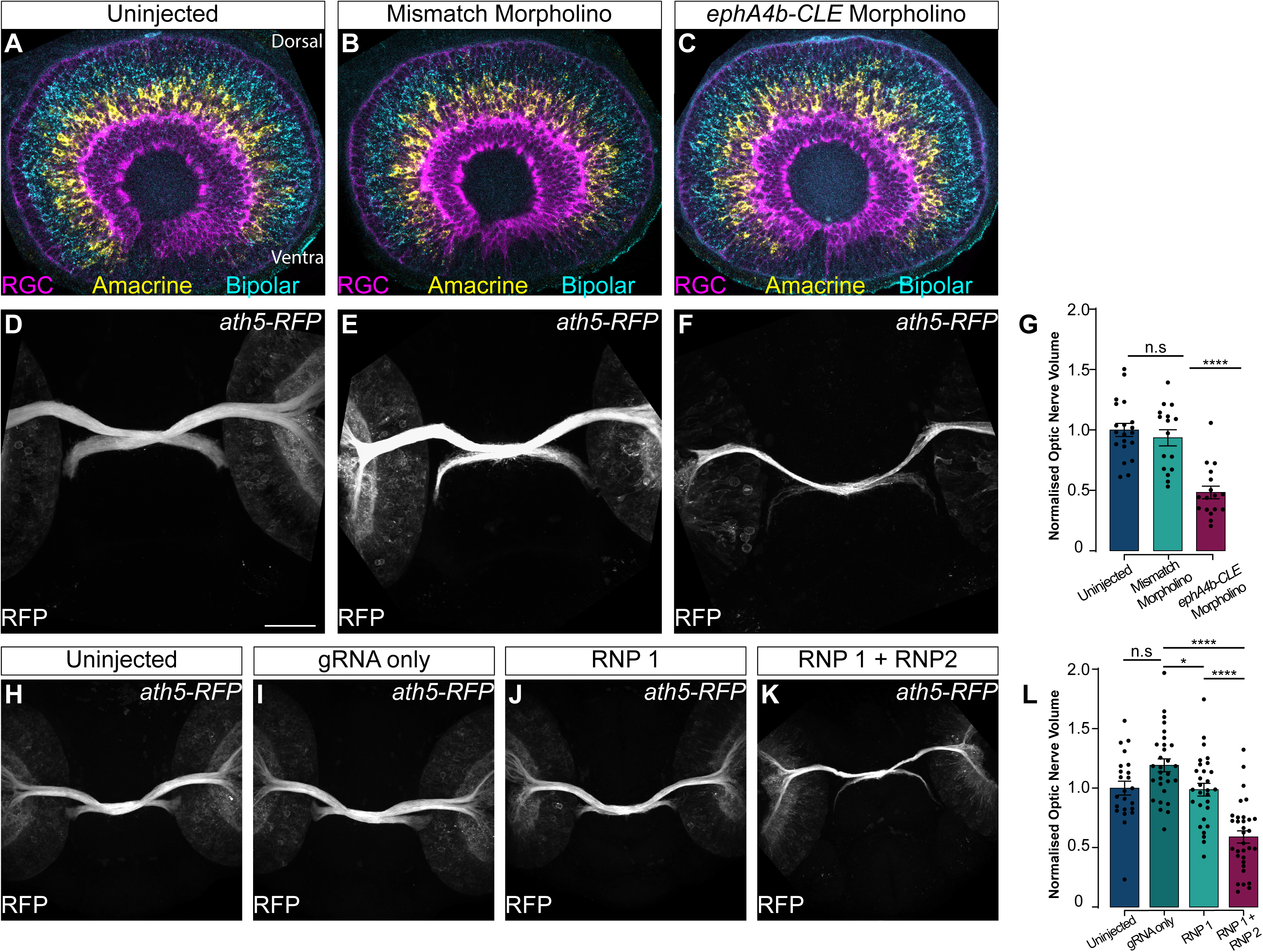
Independent disruption of ephA4b-CLE reduces optic-nerve growth without altering retinal organisation. **(A–C)** Retinal organisation in uninjected (A), mismatch morpholino-injected (B), and *ephA4b-CLE* splice-morpholino-injected (C) embryos. Retinal ganglion cells (RGCs) are shown in magenta using *tg(ath5:RFP)*, with amacrine and bipolar cell populations shown in yellow and cyan, respectively. No gross disruption of retinal organisation was observed following *ephA4b-CLE* morpholino injection. **(D–F)** Maximum-intensity projections of *tg(ath5:RFP)* optic nerves from uninjected (D), mismatch morpholino-injected (E), and *ephA4b-CLE* splice-morpholino-injected (F) embryos at 48 hpf. **(G)** Quantification of normalised optic-nerve volume. Uninjected, N=20; mismatch morpholino, N=16; *ephA4b-CLE* morpholino, N=17. Optic-nerve volume was reduced following *ephA4b-CLE* splice blocking; linear mixed-effects model, n.s., not significant; ****P<0.0001. **(H–K)** Maximum-intensity projections of 48-hpf optic nerves from uninjected (H), gRNA-only-injected (I), single-RNP-injected (J), and dual-RNP *ephA4b-CLE* F0 crispant (K) embryos. **(L)** Quantification of normalised optic-nerve volume. Uninjected, N=23; gRNA-only, N=29; RNP1, N=29; RNP1+RNP2, N=32. Statistical significance was assessed using a linear mixed-effects model; n.s., not significant; *P<0.05; ****P<0.0001. Data are shown as mean ± SD. Scale bars = 50µm.

**Supplementary Figure 8:**
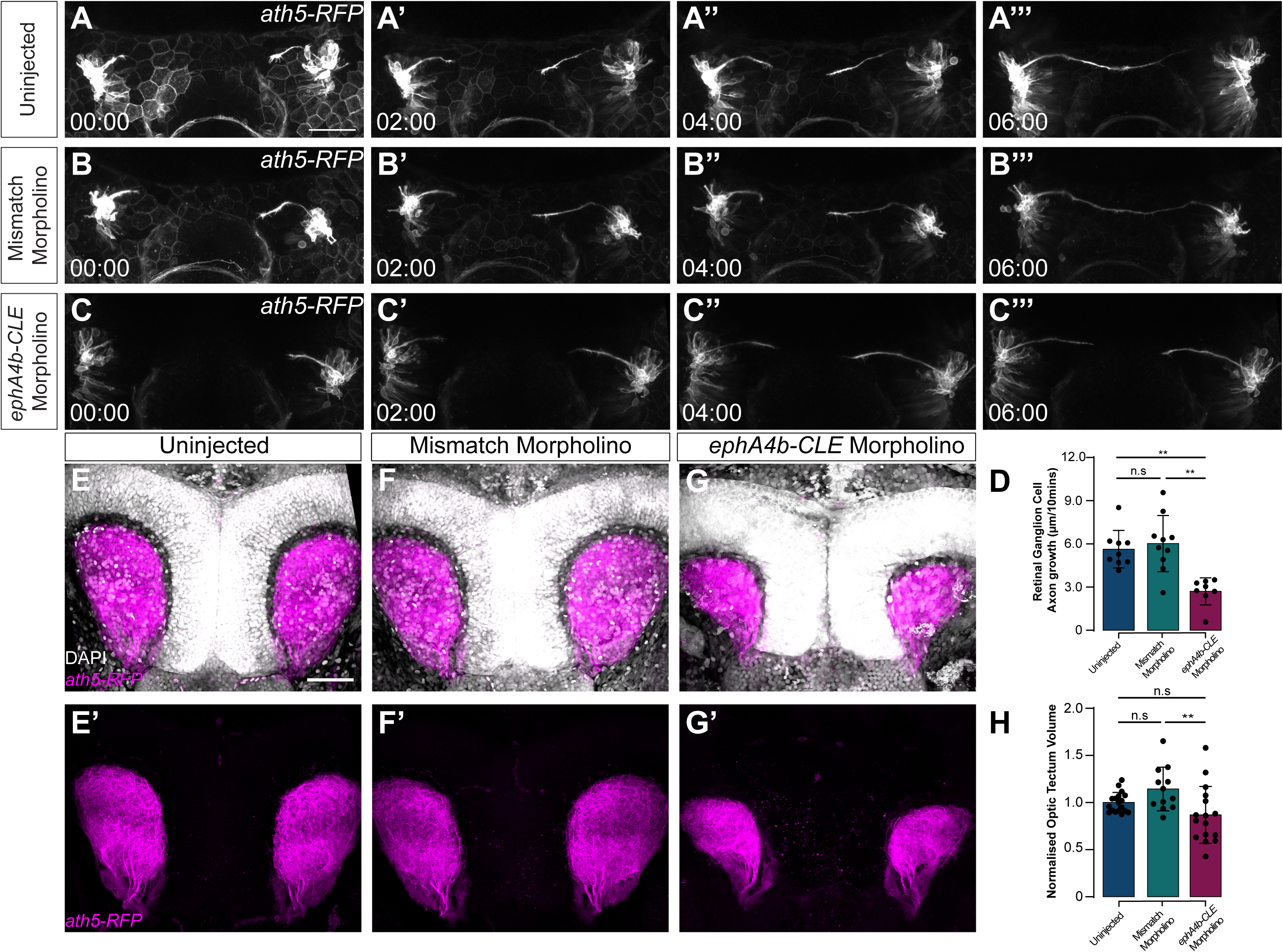
Splice-blocking loss of ephA4b-CLE reduces retinal ganglion cell axon growth and tectal projection. (A–C′′′) Time-lapse confocal imaging of pioneer RGC axon growth in uninjected (A–A′′′), mismatch morpholino-injected (B–B′′′), and *ephA4b-CLE* splice-morpholino-injected (C–C′′′) *tg(ath5:RFP)* embryos over 6 h. Images are shown at 0, 2, 4 and 6 h. **(D)** Quantification of pioneer RGC axon growth, measured as mean distance advanced per 10-min interval. Uninjected, N=9; mismatch morpholino, N=10; *ephA4b-CLE* morpholino, N=8. Statistical significance was assessed using a Kruskal–Wallis test followed by Dunn’s multiple-comparison test; n.s., not significant; **P<0.01. **(E–G)** Representative dorsal views of RGC projections into the optic tectum at 5 dpf from uninjected (E), mismatch morpholino-injected (F), and *ephA4b-CLE* morpholino-injected (G) larvae. DAPI is shown in grey and ath5:RFP-positive projections in magenta. (E′–G′) Corresponding ath5:RFP channel shown separately. **(H)** Quantification of normalised optic-tectum projection volume. *ephA4b-CLE* morpholino-injected larvae showed reduced tectal projection volume compared with controls; Kruskal–Wallis test followed by Dunn’s multiple-comparison test; n.s., not significant; **P<0.01. Data are shown as mean ± SD. Scale bars = 50µm.

**Supplementary Figure 9:**
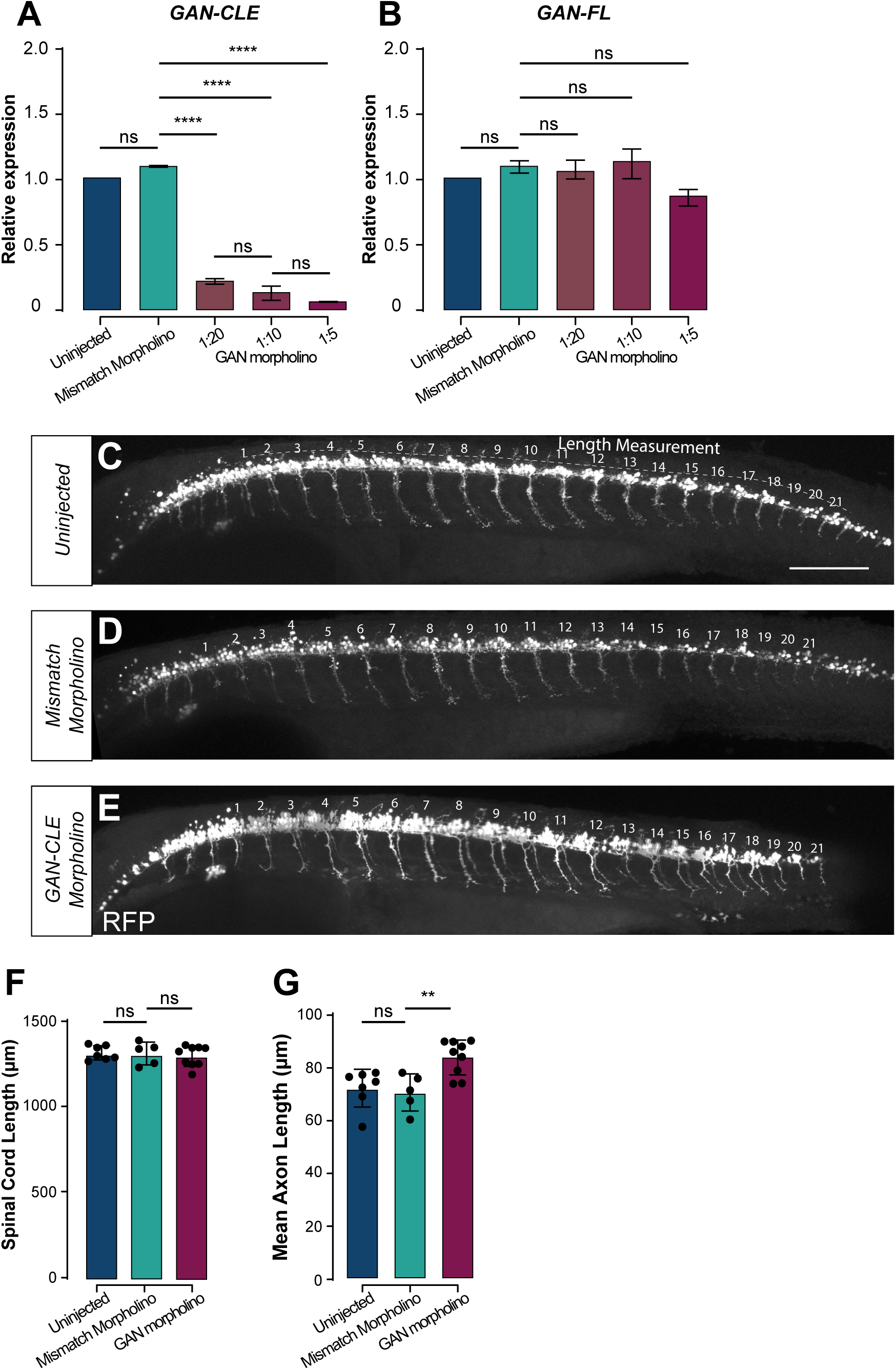
An independent splice-blocking perturbation selectively reduces GAN-CLE and recapitulates increased motor-axon length. **(A)** RT-qPCR quantification of *GAN-CLE* expression following injection of a *GAN-CLE* splice-blocking morpholino across a dilution series. *GAN-CLE* was strongly reduced relative to uninjected embryos (1.000) at 1:20 (0.196), 1:10 (0.123) and 1:5 (0.050), whereas a mismatch control morpholino had no effect (1.099). Ordinary one-way ANOVA, P=3.85×10⁻⁷; all morpholino doses versus uninjected, Tukey-adjusted P<7×10⁻⁶. **(B)** RT-qPCR quantification of full-length *GAN* transcript in the same conditions. Full-length *GAN* abundance was not significantly altered at any morpholino dose (ordinary one-way ANOVA, P=0.063), consistent with selective disruption of the CLE-containing isoform. **(C–E)** Representative lateral maximum-intensity projections of *Tg(mnx1:RFP)* embryos at 48 hpf showing spinal motor neurons and their peripheral axonal projections in uninjected (C), mismatch morpholino-injected (D) and *GAN-CLE* splice-morpholino-injected (E) embryos. Individual motor axons are numbered 1–21 along the rostrocaudal axis; the dashed line in (C) indicates the axis along which spinal-cord length was measured. (F) Quantification of spinal-cord length. Overall spinal-cord length was unchanged following *GAN-CLE* splice blocking (uninjected, 1314.9 µm, N=7; mismatch morpholino, 1311.2 µm, N=5; *GAN-CLE* morpholino, 1296.2 µm, N=9; ordinary one-way ANOVA, P=0.786). **(G)** Quantification of mean motor-axon length per embryo, averaged across the numbered axons in (C–E). Mean axon length was significantly increased in *GAN-CLE* morpholino-injected embryos (84.0 µm) compared with uninjected (72.4 µm) and mismatch morpholino-injected (70.8 µm) controls; ordinary one-way ANOVA, P=0.0029, Tukey-adjusted P=0.0097 versus uninjected and P=0.0079 versus mismatch. Individual embryo-level data points are overlaid on the summary statistics. Data are shown as mean ± SD. n.s., not significant; **P<0.01; ****P<0.0001. Scale bar = 100 µm.

## Notes

### Competing Interest Statement

The authors have declared no competing interest.

### Summary of Updates

Cryptic last exons (CLEs) are unannotated terminal exons that arise from intronic sequence, coupling intronic splice acceptors to premature stop codons and polyadenylation signals. Because they were first described in zebrafish sfpq mutants, SFPQ/TDP-43-deficient mouse cortex and ALS patient-derived neurons, CLEs have been treated almost exclusively as pathological products of RNA-binding-protein (RBP) failure. Whether cryptic terminal splicing also operates under physiological conditions has remained unresolved. Here we identify thousands of CLE-containing transcripts in wild-type zebrafish, mouse and human transcriptomes using a depth-controlled discovery pipeline that requires direct split-read support for the upstream-exon-to-CLE junction within individual libraries. CLE sequences show essentially no evolutionary constraint: median phastCons scores are indistinguishable from size-matched intronic sequence and far below conventional alternative last exons (ALEs), upstream exons and constitutive exons, and cross-species BLAST hit rates are 4- to 40-fold lower than for ALEs. CLEs are correspondingly transposon-rich, retaining the repeat character of their host introns even when compared with ALEs from the same gene. Nevertheless, CLE occurrence recurs in orthologous genes above permutation expectation and, less frequently, within orthologous introns, indicating that gene architecture creates recurrent opportunities for CLE formation without conservation of the underlying sequence. Quantifying CLE and competing full-length splicing from a shared upstream donor across stage-resolved RNA-seq, we find that CLE usage is inversely related to full-length splice usage once host-gene transcription is accounted for, and that a defined subset of CLEs is developmentally regulated (15/394 zebrafish, 6/262 mouse, 48/580 human CLEs; FDR < 0.05, beta-binomial modelling). Across 20 independent neural RBP perturbation comparisons, CLE usage is markedly more labile than ALEs matched one-to-one for baseline PSI and donor coverage (significant in 18/20 comparisons; median Cliff's δ 0.57) and than the constitutive terminal exon of the same host gene (median 84% of genes), with a subset transitioning from effectively silent to reproducibly included after perturbation. CLE-containing transcripts associate with ribosomes, CLE-specific peptides are detectable in public neural proteomes, and endogenous ALFA tagging demonstrates that ephA4b-CLE produces a stable truncated protein during normal development. Loss of ephA4b-CLE impairs retinal ganglion cell axon growth and tectal innervation, rescued dose-dependently by CLE re-expression, while disruption of GAN-CLE increases motor-axon extension and branching. Independently arising human and zebrafish EphA4 CLEs converge on similar predicted ephrin-binding ectodomains despite lacking sequence conservation. CLEs are therefore a physiological, RBP-sensitive source of transcript diversity within which individual isoforms acquire gene-specific developmental functions.

